# Gamma CV as a Marker of Circadian Disruption in C57BL/6J Mice: Correlating Neural Desynchrony with Locomotor, Thermal, and Sleep Dysrhythmia across a Spectrum of Circadian Rhythms Disruption paradigms

**DOI:** 10.64898/2026.05.01.722075

**Authors:** G. D’aloisio, A. Gekhtina, K. Laney, T. Brown, D. Moreira-Silva, A. Leake, C. Langdale, J. Gamsby, D. Gulick

## Abstract

2)

**Background:** Circadian rhythm desynchrony (CD) occurs when there is a mismatch between the circadian clock and local time, such as shift work. Mouse models are commonly employed to study CD, but may have significant shortcomings such as environmental masking, a focus only on sleep physiology, and significant variability between study designs.

**Objective:** This study used in vivo telemetry for simultaneous, real-time monitoring of locomotor activity (LA), core body temperature (CBT), and brain activity (EEG) in freely moving C57BL/6J mice to assess CD effects.

**Methods:** Four-month-old C57BL/6J mice (n=11) were surgically implanted with telemeters enabling simultaneous real-time recording of LA, CBT, EEG.: Mice were sequentially exposed to a control condition standard 12:12h light-dark cycle (T24) then 4, 8-day CD paradigms: 10:10 h short day (T20), social jet lag (SJL), repeated 6h phase advances (6A2), and a 3:3 h ultradian cycle (T6)For each paradigm, the final 48h of data (250 Hz) were analyzed.

**Results:** We found clear differences in the severity of the effects of each CD paradigm on sleep and circadian fitness, where T20∼T6>SJL>6A2. CBT revealed broader disruption, but EEG outputs proved the most sensitive indicators of internal desynchrony.

**Conclusions:** Each CD paradigm produced a unique profile across behavioral, physiological, and neural domains. We have also identified Gamma CV as a novel, sensitive metric of CD. These results highlight the necessity of multimodal monitoring to accurately characterize the impact of ecologically relevant stressors on circadian and sleep physiology.

**Statement of Significance:** Circadian rhythm desynchrony (CD), driven by shift work, jet lag, and modern irregular light exposure, is a major health burden linked to metabolic, neurodegenerative, and neuropsychiatric diseases. However, standard methods for measuring CD in laboratory models often rely on simple locomotor activity, which can “mask” the true extent of internal circadian stress. In this study, we simultaneously monitored brain EEG activity, core body temperature, and motion across four distinct models of circadian stress. We discovered that locomotor activity is a deceptive indicator of health; while mice appeared to show no alterations under several stress paradigms, their brain waves and body temperatures revealed the underlying impact of CD. Specifically, we identified “Gamma CV” as a highly sensitive new brain-wave marker that detects early circuit instability even when behavior appears normal and sleep quantity is preserved. These findings provide a marker for identifying early neurological vulnerability to irregular light schedules, offering a potential bridge to understanding similar gamma brain-wave alterations seen in addiction, early-stage Alzheimer’s disease, and other disorders.

## 3) Introduction

Circadian rhythms are near 24-hour biological cycles that exist in virtually all living organisms on the planet. This adaptive biological trait allows us to anticipate and prepare for environmental changes driven by our planet’s 24-hour rotation, such as the daily light/dark cycle. Normal circadian rhythm functionality is fundamental to human health, as a wide array of molecular, cellular, and physiological processes are influenced by the circadian regulatory system [1,2]. A key feature of human circadian rhythms is that they are reset or “entrained” by daily morning light exposure [3–6]. This temporal organization is coordinated by a central pacemaker located within the hypothalamic suprachiasmatic nucleus (SCN). The SCN consists of a network of single-cell neuronal oscillators that regulate rhythms in behavior and physiology—including locomotor activity, feeding, core body temperature, hormone secretion, and sleep-wake cycles—through direct and indirect output pathways to specific brain regions and beyond [7–9]. Thus, the SCN functions to orchestrate the synchronization of subsidiary circadian oscillators throughout the brain and in virtually all peripheral tissues, which act as local pacemakers for specific rhythmic modalities [10,11].

In modern times, humans frequently maintain schedules that oppose their natural, diurnal rhythm. This opposition creates circadian desynchrony (CD) – a mismatch of the endogenous circadian clock with external time [2,12]. As the circadian clock directly influences diverse molecular, cellular, and physiological processes, disruption of normal clock timing is an emerging health burden [1,13,14]. Indeed, CD has been shown to be either a causal or contributing factor in the pathogenesis of numerous diseases and neuropsychiatric disorders, including neurodegeneration, metabolic syndrome, mood disorders, and addiction [15–23]. Thus, there is a need to develop methodologies that measure the impact of CD on sleep and circadian fitness– particularly in the mouse, which is the most widely studied model organism. Surprisingly, while many CD protocols have been described in the literature, few studies have used more than a single protocol, making it difficult to compare findings across studies. Furthermore, there is a need to develop better methodology to assess the impact of CD, as traditional methodologies for assessing CD often rely on peripheral outputs, such as locomotor activity (LA) and core body temperature (CBT) [24,25]. LA is highly susceptible to masking: the immediate, non-circadian influence of factors such as light exposure, exercise, or other environmental stimuli [26–28]. This is further worsened by the fact that voluntary wheel running, a common method of measure LA, is itself an intervention that can improve CD in rodent models of metabolic diseases, aging, and “circadian syndrome” [29–32].

The field of circadian medicine is rapidly expanding, yet the need remains for a comprehensive, integrated approach that simultaneously monitors multiple *in vivo* outputs, including behavior (e.g. LA), physiology (e.g., CBT), and direct measures of neural circuit integrity (e.g., electroencephalogram; EEG) alongside sleep/wake cycles [33–35]. Considering the widespread use of LA and CBT as proxies for circadian status, a critical gap remains in our ability to distinguish between environmental masking and circadian fitness [26]. Specifically, we lack a clear understanding of how neural circuit integrity and sleep architecture are compromised when the circadian clock is challenged by commonly-employed CD paradigms. To begin to address this topic, have developed a novel multi-modal, *in vivo* approach with simultaneous real-time monitoring of these four integrated metrics using implanted telemetry devices.

Sleep physiology is another process that is strongly impacted by CD, yet studies investigating the specific effects of these protocols on sleep architecture remain limited. Furthermore, the acute impact of commonly employed CD paradigms on sleep and circadian fitness has largely been unexplored. For example, this gap has significant implications for understanding the effects of CD on cognitive and emotional behavior, both of which are heavily modulated by sleep quality, duration, and circadian fitness [36]. Sleep itself is a complex biological process arising from the interaction of numerous brain regions and neurotransmitter systems [37]. According to the foundational Two-Process Model, sleep is regulated by two primary mechanisms: the circadian Process C, which is centered on sleep timing and dependent on the circadian clock, and Process S, which represents sleep pressure that accumulates during wakefulness and dissipates during sleep [38,39]. These interactions are critical because changes over the sleep/wake cycle, as well as following sleep deprivation, have profound consequences for mood, synaptic plasticity, and higher-order cognition [40–42]. As sleep is central to human health, a stronger, more mechanistic understanding of how commonly employed CD paradigms acutely impact sleep is sorely needed in animal models.

In the present study, we evaluated four commonly employed CD paradigms. Two of these paradigms were based on varying ‘T cycles,’ where ‘T’ represents the period of a single light-dark cycle (e.g., T24 for a standard 24-hour day) [43]. By manipulating the T cycle length, we can evaluate how the circadian system responds to a spectrum of distinct CD stressors. Typically, strategies used to induce CD involve altering the typical daily cycle of 12 hours light followed by 12 hours dark (12:12, T24) used in rodent housing facilities. In addition, we also selected a shift work-like schedule that constantly advanced the waking and sleeping times, and a weekend-weekday-like rotating schedule. Specifically, our paradigms included: **1.** a short day of 10 hours of light and 10 hours of dark (10:10; T20), **2.** a schedule of 6-hour daily phase advances (6A2), **3.** a social jet lag (SJL) schedule of alternating between 3 days of 12:12 and 3 days of 18 hours of light and 6 hours of dark (18:6), and **4.** an ultradian T-cycle of 3 hours of light and 3 hours of dark (3:3; T6). The T20 cycle is a robust model of chronic shift work, frequently utilized for its capacity to induce CD [44].

Prior research indicates that a 10-day T20 exposure causes severe rhythm flattening, entrainment defects, and increased anxiety in several strains of mice [45,46]. T20 also alters fetal programming, where the period of pups born in DD (constant darkness) follows the maternal T20 cycle [47]. Furthermore, T20 cycles weaken SCN network coupling to the point that the mechanical and environmental stress of preparing SCN slices for ex vivo recording triggers a complete phase reset. This ‘resetting’ effect was not observed in controls, proving that the T20 paradigm reduces the integrity of the master clock.[48].

A more extreme T cycle, modeling military and emergency responder environments, we also evaluated the ultradian T6 cycle. This manipulation pushes the circadian system beyond its physiological limits of entrainment [43], effectively bypassing the master clock’s control and forcing sleep into an ultradian pattern driven primarily by the direct masking effects of light [49].

In contrast, the modest social jet lag (SJL) paradigm simulates a repetitive and significant misalignment pattern of shifting activity between weekdays and weekends—chosen for its high clinical relevance Up to 70% of the population experience at least one hour of SJL, typically later sleep on weekends, and 30–40% experience two or more hours of SJL [12,50,51].In rodents, chronic exposure to SJL generates hippocampal-dependent memory impairments, including deficits in object recognition memory and suppressed hippocampal neurogenesis promoting higher anxiety-like and depressive-like behaviors [52]. SJL also induces metabolic alterations in C57BL/6J mice, characterized by weight gain, higher fasting blood glucose, and impaired glucose tolerance [53].

Finally, we evaluated the 6A2 paradigm (6-hour phase advance every 2 days) that models the critical challenge of acute, repetitive shifts, such as in rotating shift work, a model known to cause forced desynchronization of central and peripheral rhythms [54,55]. Chronic exposure to 6A2 leads to altered metabolism [56] and also results in long-term deficits in motivation, increased anxiety, anhedonia, depressive-like behavior and metabolic alterations [55].

We hypothesized that the different CD paradigms would induce distinct, measurable patterns of misalignment and that different metrics would be differentially sensitive to circadian disruption. For instance, a study in tryptophan hydroxylase 2 null mice demonstrated that CBT rhythms were preserved, yet LA and active wakefulness were markedly altered relative to wild-type controls, leading to poor sleep–wake consolidation [57]. Consistent with this differential sensitivity of circadian outputs, LA and CBT rhythms re-entrain asynchronously following light-dark phase shifts in rats [58]. In addition, exposure to ultradian light–dark cycles such as T7 (3.5 h light/3.5 h dark), which is closely related to our T6 paradigm, has been shown to lengthen the measure of circadian period and reduce the amplitude of LA rhythms without abolishing circadian rhythmicity. Importantly, studies indicate that under such ultradian lighting conditions, the SCN clock remains functional, and core physiological rhythms, including sleep/wake organization and body temperature, do not become arrhythmic, despite marked alterations in behavioral rhythm patterns [59–62].

## 4) Material and Methods

### 4a. Subjects

Eleven C57BL/6J mice (6 males and 5 females) were received from Jackson Laboratory (Bar Harbor, ME) at 3 months of age. They were initially group-housed by sex in micro-isolation cages within the USF Health Byrd Alzheimer’s Institute vivarium. **Housing:** Standard housing conditions included a 12-hour light/dark cycle (lights on at 6:00am; ZT0) with *ad libitum* access to food and water. Mice were maintained under these standard conditions for one month. At 4 months of age, telemeter implantation (Stellar Implantable Transmitter, Type BTA-XS-C, Model #430001-IMP-LGC, FCC ID:2AC4C-AU430001LGC, TSE Systems, USA) surgery was completed (described in detail below), and mice were single housed during the post-operative recovery period. Following recovery, animals were transferred to single-housed static cages with free access to food and water inside environmental control chambers (Tecniplast Aria Bio-C36, Buguggiate, Italy). These chambers allow for the precise, real-time control of luminance and temperature which is required to program each of the CD paradigms evaluated. **Animal Care:** All experiments were carried out in accordance with the National Institutes of Health guide for the care and use of Laboratory animals (NIH Publications No. 8023, revised 2011) and were approved by the University of South Florida Institutional Animal Care and Use Committee (IACUC; PHS Assurance number: D-16-00589 [A4100-01]).

### 4b. Experimental Procedures

#### 4b1. Surgical procedure- telemeter implantation surgery. Preoperative Preparation

Two days prior to surgery, mice underwent preparatory procedures. Animals (weight range: 22–32g) were anesthetized for 10 minutes using isoflurane (1.5–2.5% in 0.5–1.5% Oxygen) and positioned on a heating pad with a nose cone. Ophthalmic veterinary ointment (Dechra Veterinary Products, KS, USA) was applied. The surgical site was shaved, prepared, and cleaned with 1X phosphate-buffered saline (PBS). To promote healthy recovery and prevent scratching of the surgical area, each mouse’s nails were trimmed. **Surgical Procedure:** On the day of surgery, pre-operative pain management was initiated with Meloxicam (Wedgewood Connect, CA, USA) 5 mg/kg, administered for 4 days. Mice were anesthetized with isoflurane (1.5-2.5% in 0.5-1.5% oxygen), and the surgical site was sterilized with Avagard (3M, MN, USA). Aseptic surgery was performed using a stereotaxic instrument (Model 940, KOPFT instruments, USA). Each mouse was positioned on a heating platform with continuous temperature monitoring during the whole procedure and Ophthalmic veterinary ointment application. The head was secured by using soft bars (Model 921 Zygoma ear cups, KOPFT). Once breathing was stabilized, a subcutaneous pocket was created between the shoulder blades to accommodate the multichannel real-time Telemetry device (Stellar Implantable Transmitter, Type BTA-XS-C, Model #430001-IMP-LGC, FCC ID:2AC4C-AU430001LGC, TSE Systems, USA). This pocket was irrigated with 1 ml of sterile PBS. The telemetry devices were prepared by trimming the leads to an appropriate length and removing the plastic insulation from the ends. **Device Placement:** A clean vertical incision was made to visualize the skull sutures and localize bregma. Specifically, two burr holes were drilled into the prefrontal cortex (PFC) (A/P: ±1.50; M/L: ±1.00 / A/P: −1.50; M/L: ±1.00). The prepared telemeter leads were formed into a V-shape bend and inserted into the burr holes, ensuring contact with the dura mater. The leads were then secured using dental cement (Lang Dental, IL,USA) and after drying, instant brain adhesive gel (LOCTITE 454, ALZET, CA,USA). The incision was closed with absorbable sutures (Vicryl 4/0, Ethicon, GA,USA) and cleaned with Avagard followed by 1X PBS. After drying, the incision was reinforced with Vetbond™ Tissue Adhesive (3M Animal Care Products, USA). **Post-Operative Care:** Following recovery from anesthesia, animals were single housed for a two-week recovery period. They were provided with specialized surgical recovery food and water (Nutra-gel and Pure-Water gel, Bio-Serv, NJ,USA) for the initial days, and pain medication (Meloxicam, 5 mg/kg) was administered as scheduled. Animals were monitored twice daily for the total duration of the recovery period.

#### 4b2. In-vivo Telemetry recording

Following a two-week recovery period, animals were individually housed in static cages with *ad libitum* access to food and water. The cages were then transferred into environmental control chambers (Tecniplast Aria Bio-C36, Buguggiate, Italy). One chamber housed the group of control animals under a standard light/dark (LD) 12:12-hour T24 cycle, and the other was used for the circadian desynchrony (CD) group. Each chamber was equipped with the TSE Stellar Telemetry System, which included one Stellar telemetry receiver and antenna (capable of collecting data from up to eight real-time implants), amplifiers, and a radio transmitter. Additionally, an integrated desktop computer utilizes Stellar Commander v2.0.0.25 and Notocord-hem software for telemetry device activation and remote, real-time data acquisition. Animals were subjected to an initial T24 baseline period followed by a sequence of four CD paradigms (T20, 6A2, SJL, and T6) (see **Figure 1** for detailed experimental timeline and CD paradigms). Due to battery life limitations, the last 48 hours of continuous recording of each paradigm were considered for data acquisition and analysis. In-vivo EEG (sampling frequency: 250 Hz), motion, and temperature were recorded continuously for 48 hours during each CD paradigm. This resulted in one EEG data output every 0.2 ms and one LA/CBT data output every second. Using the Notocord-hem software, EEG signals (including the delta (1–4 Hz), theta (4–8 Hz), alpha (8–13 Hz), beta (13–20 Hz), low gamma (20–30 Hz), and high gamma (30–50 Hz) frequency bands), motion, and temperature data were analyzed. **(see Figure 1 for experimental design).**

**Figure 1:**
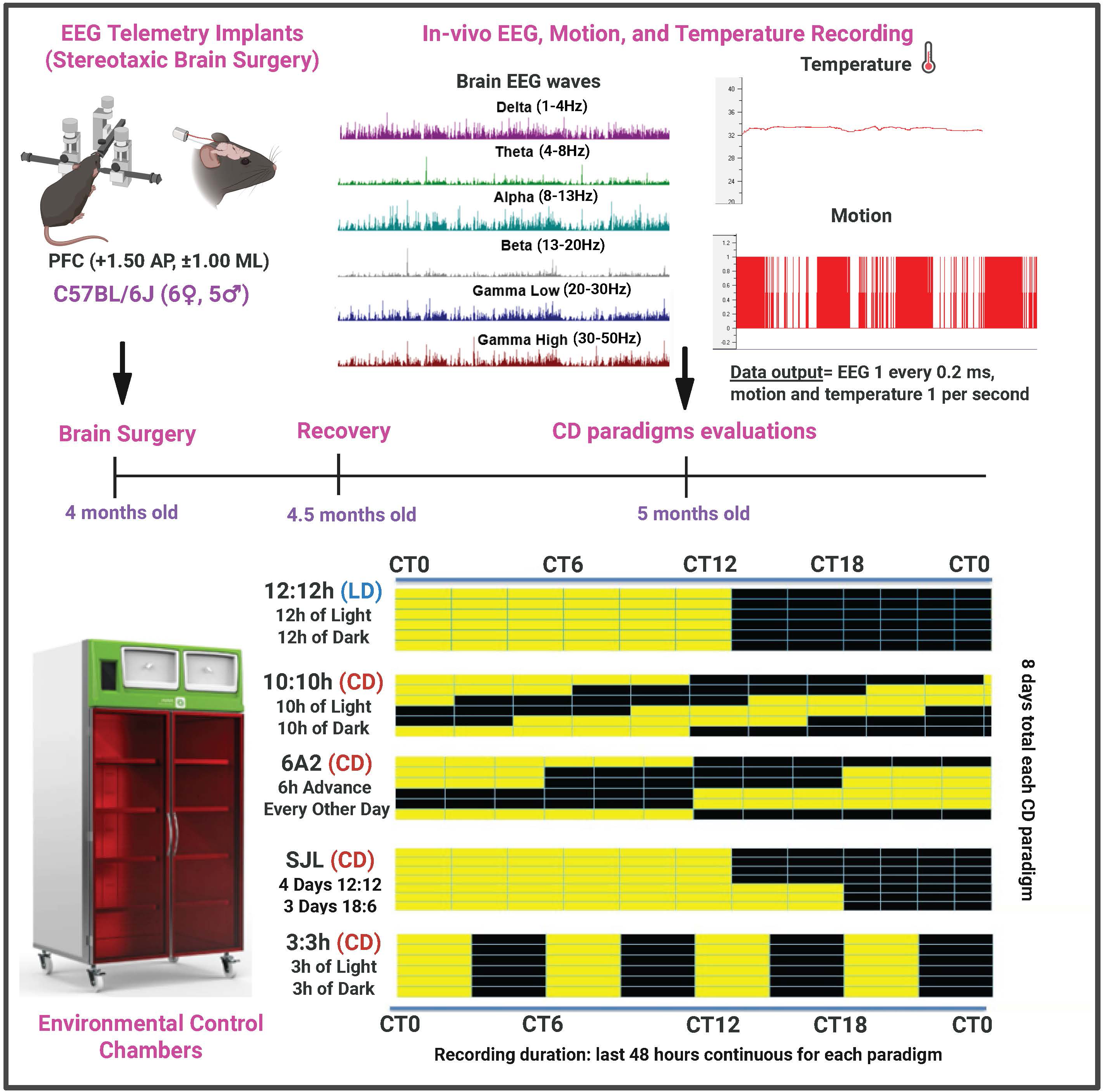
Experimental Timeline and Methodology for In-Vivo Telemetry and Circadian Disruption (CD) Paradigms. The schematic outlines the progression from surgical intervention to data acquisition. (Top Left) Stereotaxic brain surgery was performed on 4-month-old C57BL/6J mice (6 females and 5 males) to implant telemetry devices. Bipolar leads were placed in the prefrontal cortex (PFC) at coordinates AP: ±1.50; ML: ±1.00. (Top Right) Representative data traces from Notocord-hem software showing continuous in-vivo recording of brain EEG waves across six frequency bands: Delta (1–4Hz), Theta (4–8Hz), Alpha (8–13Hz), Beta (13–20Hz), Low Gamma (20–30Hz), and High Gamma (30–50Hz), alongside core body temperature (CBT) and locomotor activity (LA). Timeline and Environmental Control: Following a two-week recovery period (4.5 months old), mice were housed in Environmental Control Chambers (Bottom Left) for CD paradigm evaluations starting at 5 months of age. (Bottom Right) The experimental design included five distinct lighting paradigms (8 days each), with data acquisition focused on the final 48 hours of each: 12:12h, T24 (LD): Standard light/dark cycle (Control). 10:10h, T20 (CD): Shortened T20 cycle, 6A2 (CD): 6-hour phase advance every other day. SJL (CD): Social Jetlag simulation (4 days of 12:12; 3 days of 18:6) and 3:3h, T6 (CD): Ultradian T6 cycle. Yellow bars represent rho periods; black bars represent alpha periods. Recording resolution: EEG at 0.2 ms intervals; LA and CBT at 1 s intervals. **Alt text:** A multi-panel methodological schematic. Top panels illustrate stereotaxic telemetry implantation and representative multi-channel EEG/CBT/LA data traces. Bottom panels show the environmental housing chambers and a comparative timeline of five lighting paradigms (12:12h, 10:10h, 6A2, SJL, and 3:3h) with light/dark periods indicated by yellow and black bars.

## 5) Data collection and analysis

**Data acquisition and inspection**: In-vivo LA, CBT and EEG raw data were simultaneously collected for each telemetry device in freely moving animals. The raw data was initially imported into NOTOCORD-hem software (4.3.0.75) and underwent a visual inspection by independent, blind experts to identify potential dropouts or artifacts (e.g. abrupt spikes in power, or LA or CBT artifacts) resulting from signal interference or temporary loss of signal (in the ms-s range)**. EEG cleaning and Band separation:** Additional cleaning steps were performed on the EEG data, which was sampled every 0.2 ms, using the Operant Modules (OPR) analysis tool function within NOTOCORD-hem. 1.Offset Removal: An average value, determined by sampling stable regions of the raw EEG signal for each animal during each specific phase of evaluation, was calculated and then subtracted from the entire signal to remove any DC offset and center each sample’s average around zero. 2. Dropout Masking: The signal was converted into a binary mask of (usable data, 1s) or (dropouts or artifacts, 0s). 3. Cleaned Signal Generation: The two outputs from the offset-corrected EEG and binary mask were multiplied to produce the final cleaned EEG signal. This cleaned EEG signal was then fed into the EG10b module, which separated it into standard frequency bands defined by our research protocol: delta (1–4 Hz), theta (4–8 Hz), alpha (8–13 Hz), beta (13–20 Hz), low gamma (20–30 Hz), and high gamma (30–50 Hz). This band separation was performed using 30-second epochs to mitigate minor, transient artifacts without losing data. **LA and CBT Preprocessing:** For LA data, the RTM NOTOCORD-hem module was used for conversion into a data type suitable for exportation. For temperature, the first OPR module was used to invert and enlarge the data before restricting the output range to the specific animal’s average. The next OPR module then added this inverted and manipulated data to the raw data to adjust any dropout values to the animal’s average. **NOTOCORD-to-Excel Export:** Following the initial cleaning and processing, data from all variables were exported for subsequent post-processing using the Add-In NOTOCORD-Excel integrated tool. Raw data files for each animal were imported into Microsoft Excel using the integrated NOTOCORD Excel Add-In function to generate .xlsx data files for statistical analysis using the following settings: 1) LA and CBT data were exported and average in 30-minute values. 2) EEG data were thresholder through RMS (Root Mean Square). For each subject, the RMS cutoff value was established by visually inspecting the cleaned data alongside its associated RMS values to determine a unique threshold. This approach ensured that the analysis could exclude obviously errant data without compromising the inclusion of true physiological data points. The associated RMS values for every data point were then calculated and retained for subsequent analysis. **Clock Lab data transformation for activity profiles:** The analysis of circadian variables began by transforming the initial Motion and Temperature .xlsx data files into a format compatible with ClockLab Analysis Software (v. 6, ACTIMETRICS). These files were analyzed in the ClockLab Activity Profile to calculate Amplitude, Percent Variance (%Variance), Relative Amplitude (RA), Interdaily Stability (IS), and Intradaily Variability (IV). **Please see Figure 2**.

**Figure 2:**
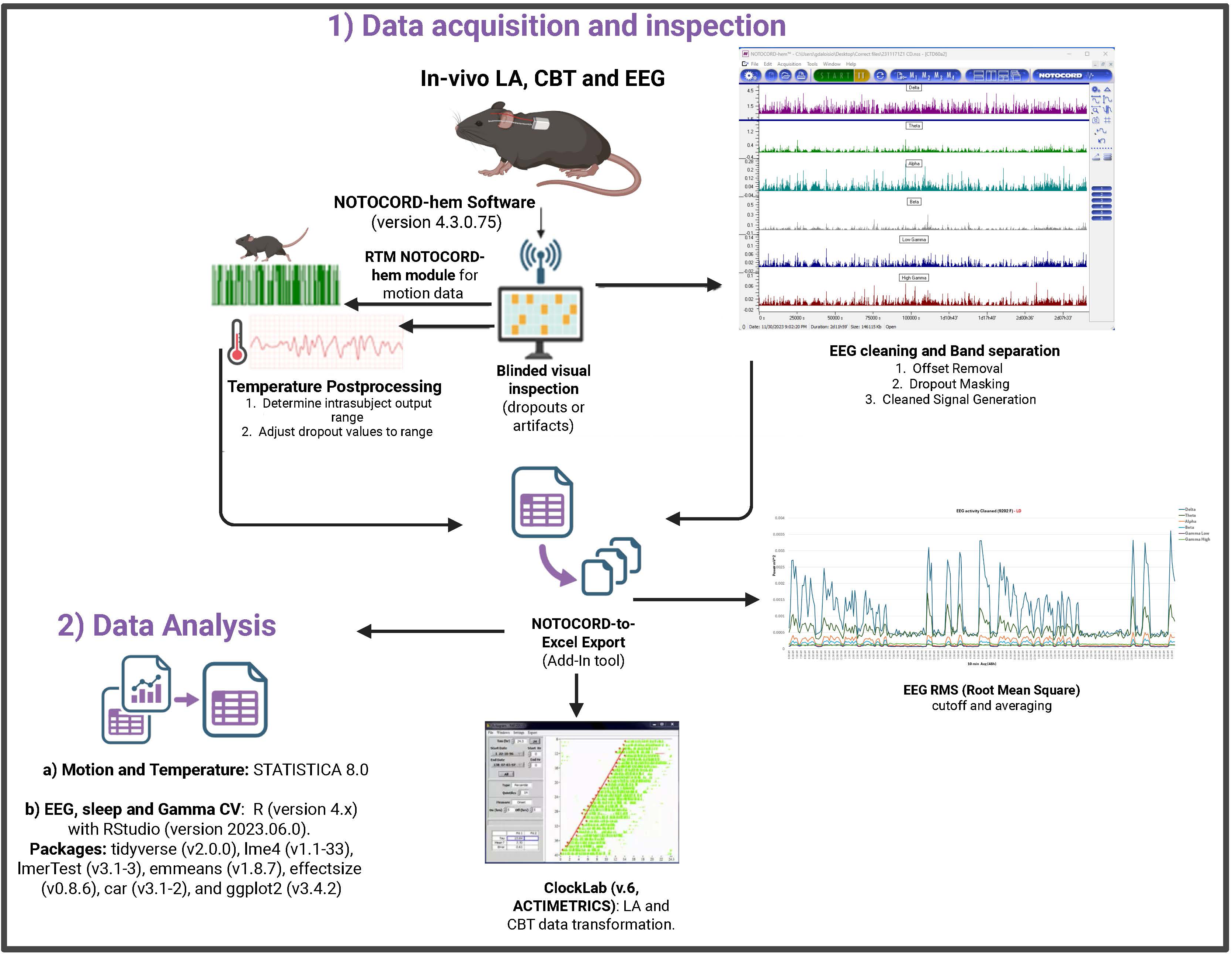
Data Acquisition, Signal Processing, and Analytical Pipeline. The flow diagram illustrates the multi-stage process performed in our high-resolution telemetry data, beginning with 1) Data Acquisition and Inspection, where raw in-vivo signals, including LA, CBT, and EEG, were recorded from freely moving mice and processed using NOTOCORD-hem software (v4.3.0.75) for blinded visual inspection of artifacts. During 2) Data Analysis, EEG data (sampled every 0.2 ms) underwent a three-stage cleaning process via Operant Modules (OPR)—including Offset Removal (OPR10a), Dropout Masking (OPR10b), and Cleaned Signal Generation (OPR10c)—to produce refined frequency bands (Delta through High Gamma) in 30-second epochs. Concurrently, LA and CBT data were preprocessed using RTM and OPR10a4/a5 modules to adjust for signal dropouts before being exported via the NOTOCORD-to-Excel Add-In for Root Mean Square (RMS) thresholding and 30-minute averaging. Final circadian rhythmicity parameters, such as Amplitude and Interdaily Stability (IS), were calculated using ClockLab (v.6). Final analysis of LA and CBT was conducted in STATISTICA 8.0, while EEG, sleep architecture, and Gamma Coefficient of Variation (CV) were analyzed using R/RStudio with specific packages (e.g. *tidyverse*, *lme4*, and *emmeans* packages). **Alt text:** A workflow diagram illustrating the multi-stage analytical pipeline for telemetry data. The process begins with raw signal acquisition (LA, CBT, EEG) in NOTOCORD-hem, followed by EEG cleaning and epoch generation through OPR modules. The pipeline shows separate processing paths for locomotor/temperature data (Excel and ClockLab) and EEG/sleep metrics (R/RStudio and STATISTICA 8.0) for final statistical modeling and rhythmicity analysis.

### Data analysis

#### 1) LA and CBT

All statistical analyses for LA and CBT data were performed using STATISTICA 8.0, encompassing the complete 48-hour recording period for each animal across all experimental evaluations. Prior to hypothesis testing, the data were rigorously screened to confirm the suitability of parametric test assumptions. Specifically, the Shapiro–Wilk test confirmed that the data exhibited a normal distribution, and Levene’s test confirmed the homogeneity of variances across the independent groups: circadian condition (LD T24 or CD) and sex (female or male). The temporal patterns of LA and CBT were analyzed using a three-way mixed-design ANOVA, with Circadian Condition (LD or CD) and Sex (female or male) as between-subject factors, and (1-hour time: 48 measures) as the within-subject factor. Additionally, phase-specific means (alpha (active/dark) and rho (rest/light) [43] of LA or CBT within each circadian paradigm were analyzed via a follow up three-way mixed ANOVA (Between factors: Circadian Condition × Sex; Within factor: Phase: light or dark). Outputs from the ClockLab analysis (e.g., Amplitude, % Variance, IS, IV, and RA were assessed using two-way ANOVAs (circadian condition × sex). Finally, any significant main effect or interaction (p<0.05) from the ANOVAs was further examined using Tukey’s Post Hoc test, with results reported as the mean ± SEM, and figures were created in GraphPad Prism (v. 10.6.1).

#### 2) EEG, sleep architecture and Gamma CV analysis

Due to the complexity of data, all statistical analyses were performed using R (version 4.x) with RStudio (version 2023.06.0). Data processing, visualization, and statistical testing utilized the following packages: tidyverse (v2.0.0), lme4 (v1.1-33), lmerTest (v3.1-3), emmeans (v1.8.7), effectsize (v0.8.6), car (v3.1-2), and ggplot2 (v3.4.2). Please refer to the supplementary results section for the precise R script used for each analysis.

##### 2.1 EEG Power Spectral Density Analysis

###### 2.1.1 Data Preprocessing and Normalization

Raw EEG power spectral density (PSD) data were extracted for six frequency bands: Delta (1-4 Hz), Theta (4-8 Hz), Alpha (8-13 Hz), Beta (13-30 Hz), Gamma Low (30-55 Hz), and Gamma High (65-100 Hz). PSD values were computed using Fast Fourier Transform (FFT) with 5-second epochs and averaged across the last 48 continuous hours of each paradigm. For each frequency band, raw power values (μV²/Hz) were Z-scored within each animal to account for inter-individual baseline differences in absolute power.

###### 2.1.2 Phase-Specific Power Analysis

For each circadian paradigm (T24, T20, 6A2, SJL,T6), EEG power was separately averaged for alpha and rho phases. To compare CD groups against the LD T24 baseline control group, we conducted separate analyses for rho phase and alpha phase comparisons during each circadian paradigm evaluation.

###### 2.1.3 Statistical Comparisons: LD Light vs CD Light Phase and LD Dark vs CD Dark Phase

For each frequency band, two-sample independent t-tests compared mean normalized PSD between the LD T24 control group (during rho phase or alpha phase) and each CD paradigm group (during rho phase or alpha phase). Statistical significance was determined using Two-sample independent t-tests by light phase, α = 0.05 with Welch’s correction for unequal variances. Multiple comparisons across paradigms were controlled using Benjamini-Hochberg False Discovery Rate (FDR) correction with q = 0.05. Significance was noted at p<0.05. Results are presented in **Figure 6A** using heatmaps with -log10(p-value) color scaling to represent statistical significance intensity. **Effect Size Calculation**: The magnitude of circadian disruption was determined by calculating Cohen’s d effect sizes for each paradigm-frequency band combination.

###### 2.1.5: Mean Power Spectral Density and Times Series

For mean power spectral density determination, each frequency band, normalized PSD (uV2/Hz) was determined separately across four experimental conditions per paradigm: LD Light (T24 control animals during rho phase), LD Dark (T24 control animals during alpha phase, CD Light (disrupted animals during rho phase), and CD Dark (disrupted animals during alpha phase). For each frequency band, a two-way ANOVA Circadian Condition (T24 control, T20, 6A2, SJL, T6) × Phase (Rho vs Alpha) was conducted on normalized power values. Type III sums of squares were used to account for unbalanced designs. When significant main effects or interactions were detected, post-hoc pairwise comparisons within each phase comparing LD vs CD were performed using estimated marginal means (emmeans package) with Tukey adjustment for multiple comparisons. To analyze the temporal dynamics differences of EEG oscillations and assess phase-specific power fluctuations, the normalized power 48h temporal series traces were generated to compare power fluctuations between the rho (orange) and alpha phase (blue) for each frequency band (Delta, Theta, Alpha, Beta, Gamma Low, and Gamma High) within the control and 4 CD paradigms (T24, T20, 6A2, SJL, T6). EEG power across all six frequency bands for each circadian paradigm, with data Z-score normalized within each animal and frequency band was considered. The Z-score transformation centers each animal’s power around zero (mean=0) with unit variance (SD=1), enabling visualization of circadian rhythm integrity (Circadian Amplitude) independent of absolute power magnitude. Statistical comparisons employed one-way ANOVA comparing circadian amplitudes across paradigms, followed by Tukey’s HSD post-hoc tests. As a follow-up step, **cross-frequency synchrony analysis was performed** to assess whether circadian rhythms remain synchronized across frequency bands, we computed mean pairwise Pearson correlations between all six frequency bands (15 unique pairs) for each paradigm’s 48-hour temporal profile. Effect sizes (Cohen’s d) were calculated as the difference in means divided by the pooled standard deviation.

##### 2.2 Sleep Architecture Analysis

###### 2.2.1 Sleep State Classification, Quantification and Analysis

Sleep-wake states were automatically scored in 10-second epochs. This process is typically based on the combined analysis of EEG power spectral characteristics and electromyographic (EMG) activity. Given the limitation that the Telemetry channel configuration did not allow for simultaneous EMG recording, we utilized the LA (activity) signal recorded by the device as a reliable proxy for muscle tone. This substitution was supported empirically by a strong positive correlation between LA and EEG-defined wakefulness (pooled Pearson r = 0.97, p < 0.001; Supplementary Figure 1). Three states were classified: Wake, Non-Rapid Eye Movement (NREM) sleep, and Rapid Eye Movement (REM) sleep.

For each 10-second epoch, states were assigned based on:

- Wake: High motion amplitude, low Delta power, high Beta/Gamma power
- NREM: Low motion amplitude, high Delta power (>70% of total power)
- REM: Low motion amplitude, high Theta power, Theta/Delta ratio > 1.5

**For each sleep state (Wake, NREM, REM),** a two-way mixed ANOVA was conducted with Phase (Rho vs Alpha; within-subjects factor) and Circadian Condition (LD vs CD; between-subjects factor) as independent variables: Statistical significance was determined at α = 0.05. Effect sizes (partial eta-squared, η²_p) were calculated using the effect size package. F-statistics, degrees of freedom, p-values, and η²_p values were calculated. When significant interactions (Phase × Circadian Condition) were observed, pairwise comparisons were conducted using Tukey’s Honest Significant Difference (HSD) test via the emmeans package. Tukey’s HSD controls the family-wise error rate across all pairwise comparisons. The following comparisons were evaluated: within LD group: LD Rho vs LD Alpha, within CD group: CD Rho vs CD Alpha, within Light phase: LD Rho vs CD Rho and within Dark phase: LD Alpha vs CD Alpha. Adjusted p-values and mean differences are reported for significant comparisons. Mean differences are expressed as percentage point differences.

**To assess the preservation or loss of light-phase discrimination in terms of sleep architecture within each experimental group,** paired t-tests were conducted comparing rho phase vs alpha phase percentages within the LD control group and within each CD group separately. Two-tailed tests were used with α = 0.05. Results are reported with t-statistics, degrees of freedom, p-values, and Cohen’s d values.

**Sleep efficiency** was calculated as the percentage of time spent in sleep (NREM + REM) out of total recording time during each phase. Two-way ANOVAs (Phase × Circadian Condition) were conducted for sleep efficiency using identical procedures as described for each sleep stage. Across all paradigms condition-specific error bars represent SEM.

To quantify the balance between NREM and REM sleep, **NREM: REM ratios were calculated for each phase**. T-statistics, degrees of freedom, p-values, and Cohen’s d effect sizes were calculated.

In an additional step of analysis, **Paradoxical Sleep-Wake activity** was analyzed. Paradoxical Activity quantifies inappropriate wake during the rho rest phase or inappropriate sleep during the alpha active phase for rodents. It was calculated as the deviation from the LD T24 baseline: **Paradoxical Activity (%) = (CD_Light_Wake - LD_Light_Wake)** where positive values indicate excessive wake during rho phase (paradoxical wakefulness) and negative values indicate excessive sleep during alpha phase (paradoxical hypersomnia). One sample t-tests evaluated whether Paradoxical Activity significantly differed from zero. Results are presented with t-statistics, degrees of freedom, p-values, and Cohen’s d values (compared to zero baseline).

Finally, to analyze the **degree of sleep architecture disruption and circadian misalignment** across sleep states (Wake, NREM, REM), phases (rho or alpha), and paradigms (T20, SJL, 6A2 and T6), percent changes from LD T24 baseline were calculated and displayed as heatmaps. Color intensity represents magnitude of misalignment (percentage point deviation), facilitating cross-paradigm comparisons.

##### 2.3 Gamma Coefficient of Variation (CV) Analysis

Gamma CV as the quantification of the temporal variability of gamma band power was calculated as a biomarker of neural circuit stability for each animal and paradigm across the total 48-hour of continuous recording using a combined gamma band power (30-100Hz): Gamma CV = SD_Gamma / Mean_Gamma. where SD_Gamma is the standard deviation of gamma power across all 10-second epochs within a phase (rho or alpha) and Mean_Gamma is the mean gamma power across those epochs. Lower CV values indicated stable, rhythmic gamma oscillations characteristic of healthy circadian regulation; higher CV values indicated unstable, arrhythmic gamma activity reflecting circadian disruption. **To test the hypothesis that Gamma CV reflects circadian amplitude preservation**, Pearson correlation analysis was conducted between Gamma CV and circadian amplitude percentage (derived from LA/CBT ClockLab analysis). Statistical significance was assessed using a two-tailed t-test. Regression coefficients (β_₀_, β_₁_), t-statistics, p-values, 95% confidence intervals, and R² are reported.

**Partial Correlation Analysis: Gamma CV vs Sleep Quantity:** To dissociate EEG quality (Gamma CV) from sleep quantity (total sleep %), partial correlation analysis was conducted, controlling circadian amplitude. This analysis tests whether Gamma CV remains associated with sleep quantity after accounting for the shared variance explained by circadian amplitude. Results, including partial correlation coefficients, t-statistics, p-values, and degrees of freedom, are reported.

**EEG Spectral Disruption Across Paradigms**: Effect Size Analysis were conducted to quantify the magnitude of EEG disruption across all frequency bands and paradigms, one-way ANOVAs were conducted for each frequency band (Delta, Theta, Alpha, Beta, Gamma Low, Gamma High) with Paradigm as the independent variable. F-statistics, degrees of freedom, and p-values are reported for each frequency band. Cohen’s d effect sizes comparing each CD paradigm to a T24 baseline were calculated and displayed in **Figure 8D** as a heatmap with color intensity representing effect size magnitude.

**Neural Quality Categories:** Based on Gamma CV thresholds, paradigms were categorized into neural quality classifications: Optimal (CV ≤ 0.30): T24 baseline, constrained (CV < 0.25): 6A2 forced adaptation, compensated (0.30 < CV ≤ 0.40): SJL phase misalignment, moderate disruption (0.40 < CV ≤ 0.60): T20 rhythm flattening, catastrophic disruption (CV > 0.60): T6 complete desynchronization. This classification framework integrates Gamma CV, circadian amplitude, sleep efficiency, and EEG spectral data to provide a unified interpretation of circadian-neural impairment across paradigms.

## 6) Results

**Main effect of sex for LA and CBT:** A significant main effect of sex was observed for both LA and CBT. This effect was consistent across all comparisons (p-values 0.000-0.049; power (1-β) 0.53-0.99) and showed no significant interaction with circadian paradigm (LD or CD) or phase (rho or alpha). Females exhibited higher levels of LA and CBT compared to males (**supplementary Figure 2**). This finding, wherein females display higher LA and CBT, is consistent with established principles of sexual dimorphism in circadian physiology, including general metabolic differences, higher heart rate and higher LA and CBT rhythms in females vs males, as reported in both human and animal studies [63–66]. Crucially, as confirmed by the absence of significant circadian condition x sex interactions, sex differences did not account for the primary effects of the circadian paradigms on any metric (LA, CBT, power spectrum density, sleep, or Gamma CV). Moreover, no significant main effect of sex nor interaction of sex with other variables was observed for EEG measures.

### CD Effects on LA

Figure 3 illustrates the temporal patterns of LA as a function of CD paradigm and rho/alpha phases. As expected in T24, since both groups were maintained under an identical 12:12h light/dark cycle, a repeated measures ANOVA (rmANOVA) across 48h confirmed no significant interaction between the circadian condition and bins (Figure 3A; F(48,288)=0.79, p=0.834). This initial analysis also established a significant main effect for bins (F(48,288)=6.08, p<0.001) and sex (F(1,7)=25.87, p=0.002, observed power= 0.99), but no effect of group, indicating that LA changed significantly over time and by sex. Further supporting the lack of group difference, a three-way ANOVA focusing on the phases confirmed that the activity profile was independent of the assigned circadian condition (non-significant phase × circadian condition interaction; F(1,7)=1.43, p=0.271). Critically, this analysis revealed a highly significant main effect of phase (F(1,7)=254.69, p<0.001), demonstrating that LA was overwhelmingly concentrated during the alpha phase compared to rho phase, as expected (Figure 3A). The main effect of sex was also confirmed in this analysis (F(1,7)=17.51, p=0.004, observed power=0.94).

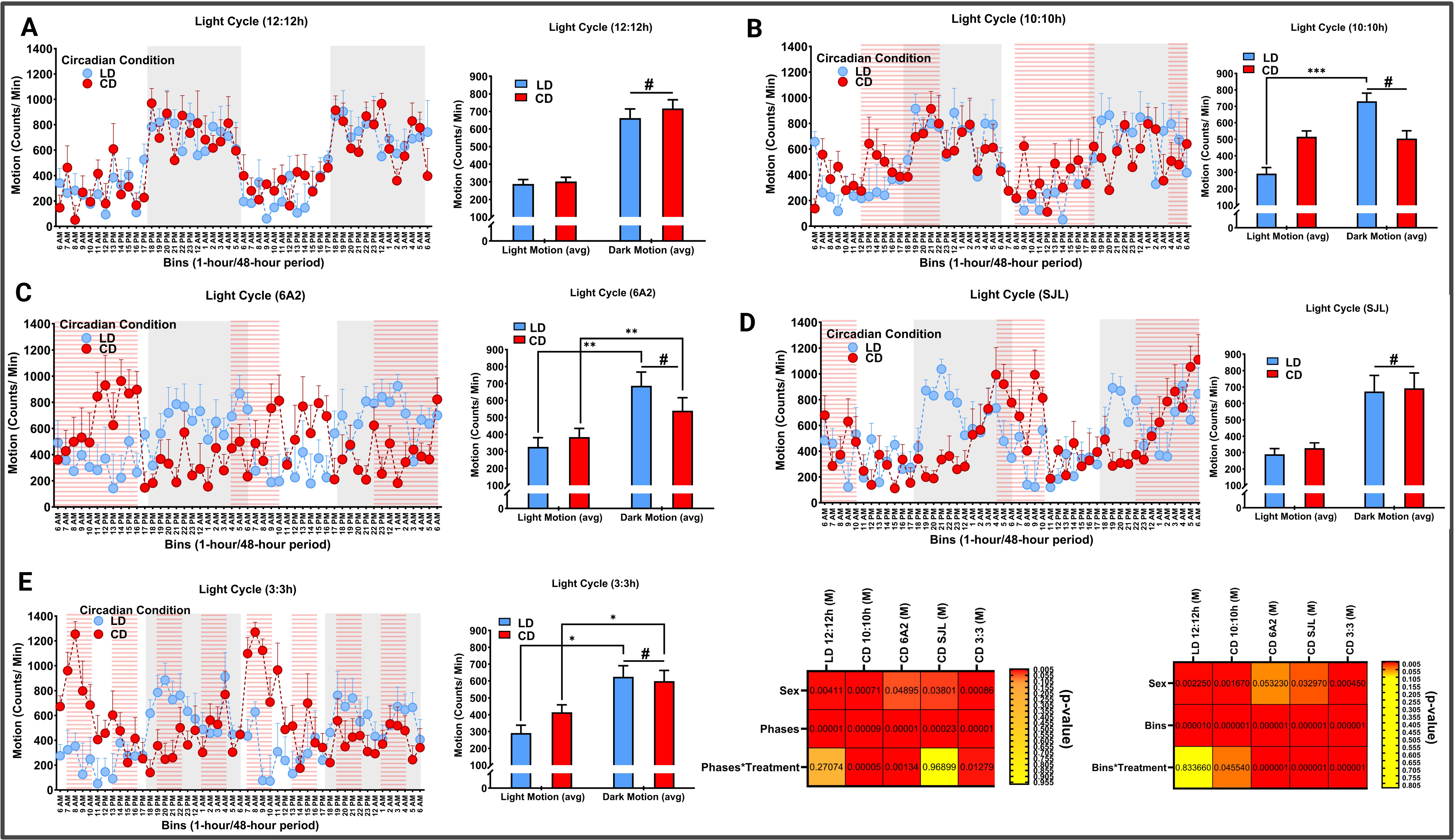

**T20:** A rmANOVA showed a main effect of bins (F(48, 336)=3.9497, p<.0001), sex (F(1, 7)=24.450, p=.00167, observed power=0.98) and a significant interaction between bins and circadian condition (F(48, 336)=1.4087, p=.04554). A follow-up three-way ANOVA focusing on rho/alpha phases showed main effects of phase (F(1, 7)=63.347, p<0.001), sex (F(1, 7)=32.925, p=.00071, observed power= 0.99) and in this case, a strong significant interaction between phase and circadian condition (F(1, 7)=76.562, p<0.001). Tukey’s comparisons indicated that only the LD T24 control animals showed higher activity during the alpha phase and lower activity during the rho phase, while similar levels of LA were observed for the group of animals under T20 regardless of light phase, a canonical sign of circadian disruption (Figure 3B).

**6A2:** An interaction of bins x circadian condition (F(48, 336)=4.1114, p<.001) was observed. In this case, the follow-up ANOVA showed a main effect of phase (F(1, 7)=167.84, p<.001), sex (F(1, 7)=5.6596, p=.04895, observed power=0.53) and a significant interaction of phase x circadian condition (F(1, 7)=26.403, p=.00134). Tukey’s post hoc tests revealed that both the T24 group and the 6A2 group exhibited significantly higher levels of LA during the alpha phase compared to their respective rho phase (Figure 3C).

**SJL:** RmANOVA showed main effects of bins (F(48, 336)=4.2370, p<.001), sex (F(1, 7)=7.0197, p=.03297, observed power=0.63) and a significant interaction between bins and circadian condition (F(48, 336)=3.4006, p<.001). Follow-up ANOVA showed main effects of sex (F(1, 7)=6.6365, p=.03801, observed power= 0.59), and phase (F(1, 7)=50.321, p<.001) but no significant interaction between phases and circadian condition, F(1, 7)=.00162, p=.96899. Animals under SJL showed a similar profile of LA as T24 controls, indicating minimal circadian disruption (Figure 3D).

**T6:** RmANOVA showed main effects for sex (F(1,7)=38.253, p<0.001, observed power=0.99) and bins (F(48, 336)=2.6293, p<.001). A significant interaction was also found between bins × circadian condition (F(48, 336)=4.6537, p<0.001). A follow-up three-way mixed measures ANOVA specifically for light phase LA revealed main effects of sex (F(1,7)=30.793, p<0.001, observed power=0.99) and phase (F(1,7)=124.39, p<0.001), as well as a significant interaction between phase × circadian condition (F(1,7)=11.012, p=0.012).Tukey’s post hoc comparisons showed that, even when exposed to an aberrant ultradian cycle, both the T24 and T6 groups exhibited significantly higher levels of LA during the alpha phase compared to their respective rho phase (within-group comparison). This result could be driven by the high peak levels of LA observed in the T6 group. Although the T6 group generally showed similar overall levels of LA independent of phase, these specific LA peaks match the ZT0 for a regular T24 cycle (Figure 3E).

### CBT

Figure 4 illustrates the temporal patterns and rho/alpha phase averages of CBT across CD paradigms. As observed in Figure 4A, during the baseline T24 cycle, the rmANOVA showed a strong main effect of bins (F(48,366)=18.496, p<0.001). However, confirming the LA analysis, and as expected for baseline synchrony, the interaction between bins and circadian condition was not significant (F(48,366)=1.4032, p=0.09869). A follow-up ANOVA for rho/alpha phase confirmed this high degree of synchrony, yielding only a main effect of phase (F(1,7)=73.811, p<.001). Both groups exhibited higher CBT during the alpha phase and lower levels during the rho phase, with no significant interaction between phase and circadian condition (F(1,7)=0.24290, p=0.63720).

**Figure 4:**
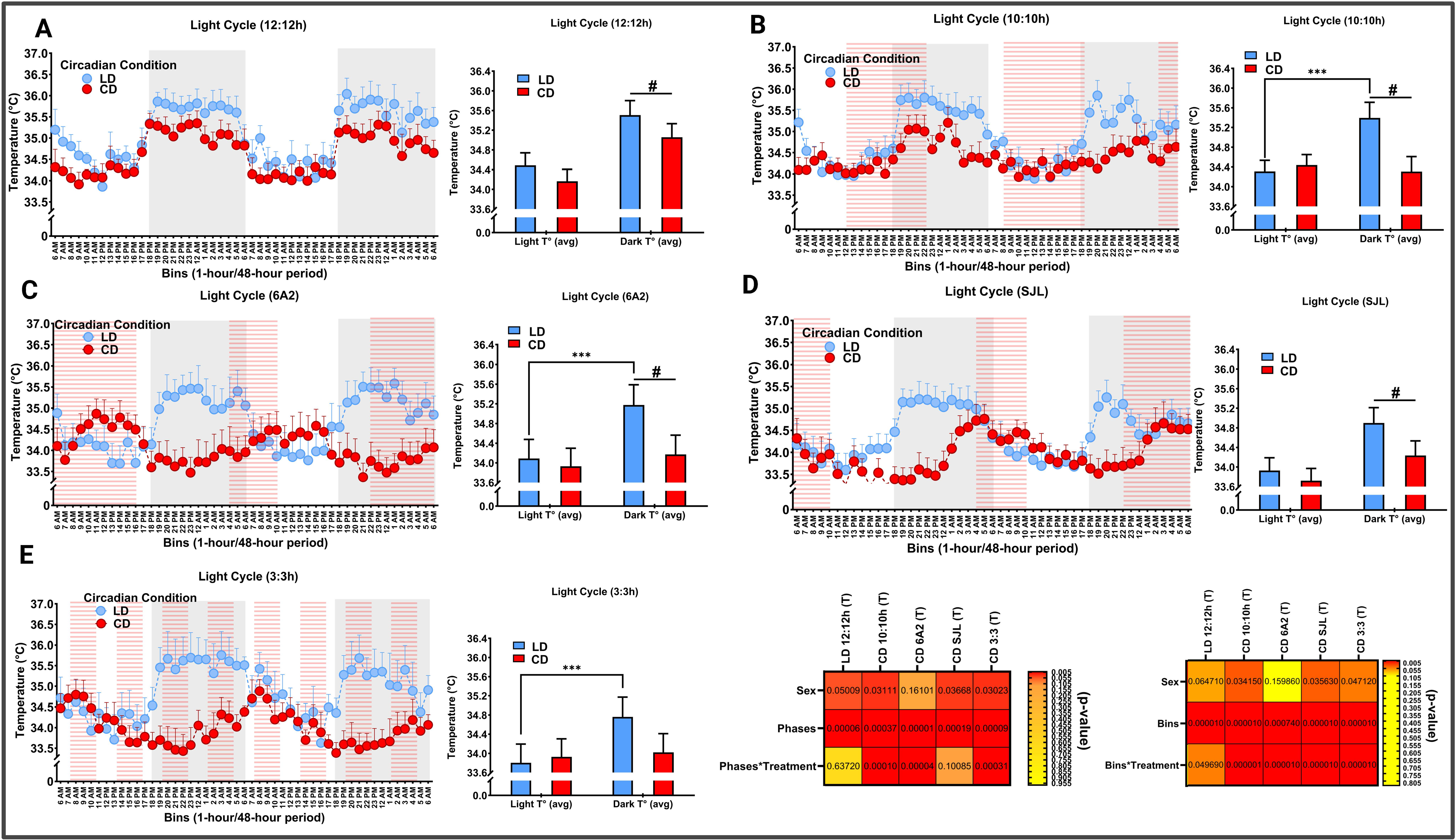
Core Body Temperature (CBT) Profiles and Circadian Misalignment Across CD Paradigms within rho/alpha Phases. The figure illustrates the temporal biothermal patterns and average CBT across standard T24 control and various CD paradigms. (A–E) Each panel presents a 48-hour CBT profile (left), where gray shaded areas represent the alpha phase for T24 controls and red hatched areas indicate the alpha phase for CD groups within each specific paradigm. Corresponding bar graphs (right) compare average CBT during the rho (light) and alpha (dark) phases. (A) T24 (LD): Shows high baseline synchrony, with both groups exhibiting significantly elevated CBT during the alpha phase (#, p < 0.001). (B) T20 (CD): Evidence of significant biothermal disruption; while T24 controls maintain rhythmicity, the T20 group exhibits statistically similar CBT levels across both phases. (C) 6A2 (CD): Despite preserved LA rhythmicity (see Fig. 3), CBT shows profound disruption with a loss of phase-dependent temperature differentiation under 6A2. (D) SJL (CD): Represents the lowest level of disruption, with CD profiles remaining statistically indistinguishable from T24 controls. (E) T6 (CD): Exposure to the ultradian cycle results in a loss of the expected rho/alpha CBT rhythm, further identifying CBT as a sensitive indicator of circadian misalignment. Heatmaps (Bottom Right): Summarize p-values for main effects (Sex, Bins, Phase) and interactions, highlighting the significant impact of CD paradigms on biothermal rhythmicity. Data are presented as mean ± SEM; symbols denote significance levels: ***p < 0.001 (inter-group, CD vs. LD), #p < 0.001 (intra-group phase difference). **Alt Text:** A multi-panel data figure displaying 48-hour CBT time-series profiles and corresponding phase-specific bar graphs across five lighting paradigms. Gray shading and red hatching indicate dark phases. A statistical heatmap in the bottom right summarizes main effects and interactions for sex, time bins, and phase across all experimental groups.

**T20:** We observed a biothermal profile that suggested a failure of circadian entrainment, aligning with the observed circadian disruption in LA. The rmANOVA across bins yielded significant main effects for sex (F(1,7)=6.8921, p=0.034, observed power=0.61) and bins (F(48, 336)=13.169, p<0.001), along with a strong significant interaction between bins × circadian condition (F(48, 336)=3.9694, p<.001). A follow-up ANOVA focused on rho/alpha phase confirmed this disruption with a strong phase × circadian condition interaction (F(1,7)=62.768, p=0.0001) and main effects for sex (F(1,7)=7.2330, p=0.031, observed power=0.63) and phase (F(1,7)=40.878, p<0.001). As visually represented in Figure 4B and confirmed by Tukey’s post hoc tests, only the T24 control group maintained the expected rhythm of significantly higher CBT during the alpha phase, whereas the T20 group exhibited the same CBT levels regardless of the current light phase, providing evidence of circadian misalignment in biothermal regulation.

**6A2:** CBT data (Figure 4C) showed a strong sign of circadian disruption, despite LA being preserved. The rmANOVA yielded a significant main effect of time (F(48, 336)=1.8779, p<.001) and a strong significant bin × circadian condition interaction (F(48, 336)=12.177, p<0.001). This was further supported by the phase analysis ANOVA, which showed a significant phase × circadian condition interaction (F(1,7)=83.612, p<0.001) and a highly significant main effect of phase (F(1,7)=205.80, p<0.001). Tukey’s post hoc comparisons revealed that only the T24 control group maintained the expected CBT rhythm; conversely, the 6A2 group exhibited the same average CBT level across both phases. Consistent with established models of circadian rhythmicity (Dijk & Czeisler, 1995), this dissociation from the LA results suggests that CBT serves as a more sensitive physiological indicator of subtle circadian disruption under our specific experimental conditions.

**SJL:** CBT results mirrored the findings for LA. The rmANOVA across bins showed significant main effects for sex (F(1,7)=6.780,p=0.03523, observed power= 0.62) and bins (F(48, 336)=7.0509, p<0.001), alongside a significant bin × circadian condition interaction (F(48, 336)=5.8666, p<0.001). Crucially, the follow-up phase ANOVA showed no significant interaction between phase × circadian condition (F(1,7)=3.568, p=0.101), although main effects for sex (F(1,7)=6.637, p=0.037, observed power= 0.60) and phase (F(1,7)=50.321, p<0.001) were observed. Tukey’s post hoc tests confirmed that the SJL group exhibited rho/alpha CBT profiles statistically indistinguishable from controls animals under T24, indicating that this paradigm resulted in the lowest level of circadian disruption (Figure 4D).

**T6:** RmANOVA showed main effects of sex (F(1, 7)=6.2036, p=.04712, observed power=0.53), bins (F(48, 336)=4.1979, p<.001) and a strong interaction between bins and circadian condition (F(48, 336)=8.5418, p<0.001). The follow-up ANOVA yielded main effects of sex (F(1, 7)=7.3408, p=.03023, observed power= 0.64), phases F(1, 7)=64.563, p<.001 and a strong interaction of phase x circadian condition F(1, 7)=43.276, p<.001). Tukey comparisons indicated that T6 showed the same levels of CBT regardless of rho/alpha phase as a sign of circadian disruption even when LA didn’t show this effect (Figure 4E).

### Circadian Fitness

A two-way ANOVAs (circadian condition × sex) revealed that no significant sex differences were observed in any circadian metric across all paradigms, a result contrasting with our LA and CBT findings. However, among all tested CD conditions (T20, SJL, 6A2, T6), only the T20 CD paradigm impaired the core circadian metrics, with no detectable deficit under other CD paradigms.

**T20:** There were significant main effects of circadian condition across circadian metrics of fitness for both LA and CBT data. The T20 group exhibited a profound deterioration in the LA rhythm, yielding significant main effects consistent with a circadian impairment: lower amplitude (F(1,9)=5.9591, p=.03730), and relative amplitude (RA; (F(1,9)=19.486, p=.00169). Measures of rhythm instability were also significantly affected: lower % variance (F(1,9)=36.900, p=.00018), reflecting reduced power, higher IV (F(1,9)=16.590, p=.00279), signaling greater rhythm fragmentation. Finally, IS was significantly lower (F(1,9)=5.4613, p=.04424), indicating inconsistency in the daily rhythm. The analysis of CBT further supported this widespread desynchronizing effect of the T20 cycle, with similar main effects: IV was markedly higher (F(1,9)=52.808, p<.001), but RA (F(1,9)=28.295, p<.0001), amplitude (F(1,9)=20.459, p=.00144), and % Variance were all lower (F(1,9)=14.005, p=.00461). A strong trend was also observed for lower IS compared with the T24 control (F(1,9)=4.4763, p=.06348). Collectively, these results suggest that the T20 cycle imposes a uniquely powerful and systemic desynchronizing stressor on the endogenous circadian clock in numerous metrics of circadian fitness (Figure 5).

**Figure 5:**
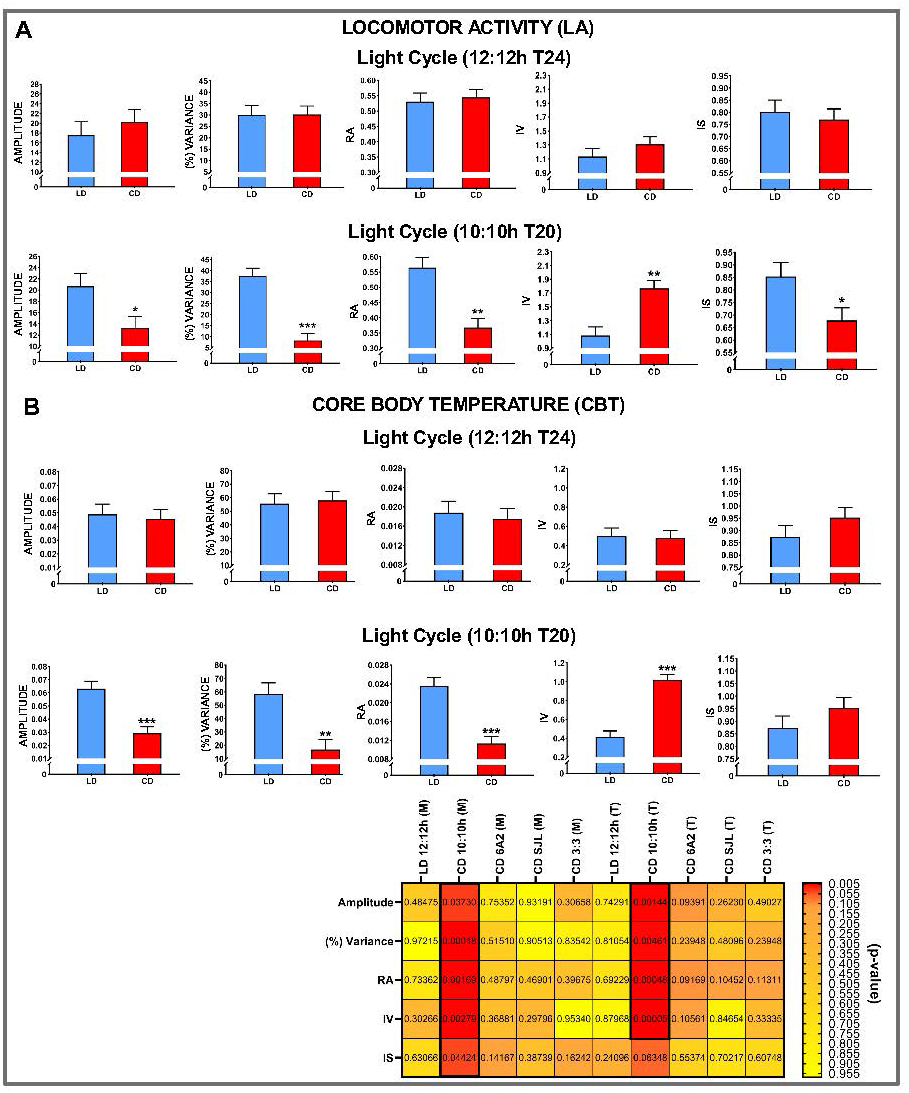
Circadian Fitness Metrics for (A) Locomotor Activity (LA) and (B) Core Body Temperature (CBT). The figure represents the impact of various CD paradigms on the robustness and stability of endogenous rhythms using five key circadian metrics: Amplitude, Percent Variance (% Variance), Relative Amplitude (RA), Intradaily Variability (IV), and Interdaily Stability (IS). Bar graphs (top and middle) illustrate results for Motion (LA) and Temperature (CBT) under standard T24 (12:12h LD) and T20 (10:10h CD) conditions. (Top and Middle Panels): While no significant sex differences were observed across paradigms, the T20 cycle uniquely and severely impaired circadian fitness. This is characterized by a profound reduction in rhythm power (lower Amplitude, RA, and % Variance) and increased rhythm fragmentation (higher IV) for both LA and CBT. Additionally, a significant decrease in IS for LA indicates reduced day-to-day consistency of the activity rhythm under T20. (Bottom Panel): The summary heatmap provides a comprehensive comparison of p-values across all CD paradigms (T20, 6A2, SJL, and T6). The distinct red column for the T20 condition highlights its status as a uniquely powerful systemic stressor, whereas other paradigms (6A2, SJL, T6) showed no detectable deficits in these specific core circadian metrics. Data are presented as mean ± SEM; symbols denote significance levels: *p < 0.05, **p < 0.01, ***p < 0.001. **Alt Text:** A multi-panel figure displaying circadian fitness metrics for locomotor activity and core body temperature. Top and middle sections feature bar graphs for five metrics (Amplitude, % Variance, RA, IV, and IS) comparing T24 and T20 cycles. A summary heatmap at the bottom compares p-values across all paradigms, with a prominent red column highlighting the systemic disruption caused uniquely by the T20 condition.

### EEG

A comprehensive characterization of brain oscillatory activity across CD paradigms revealed frequency-specific vulnerabilities to each experimental condition. During the baseline period, there were no significant differences between LD rho and CD rho phases across any frequency band, confirming baseline stability (Figure 6A).

**Figure 6:**
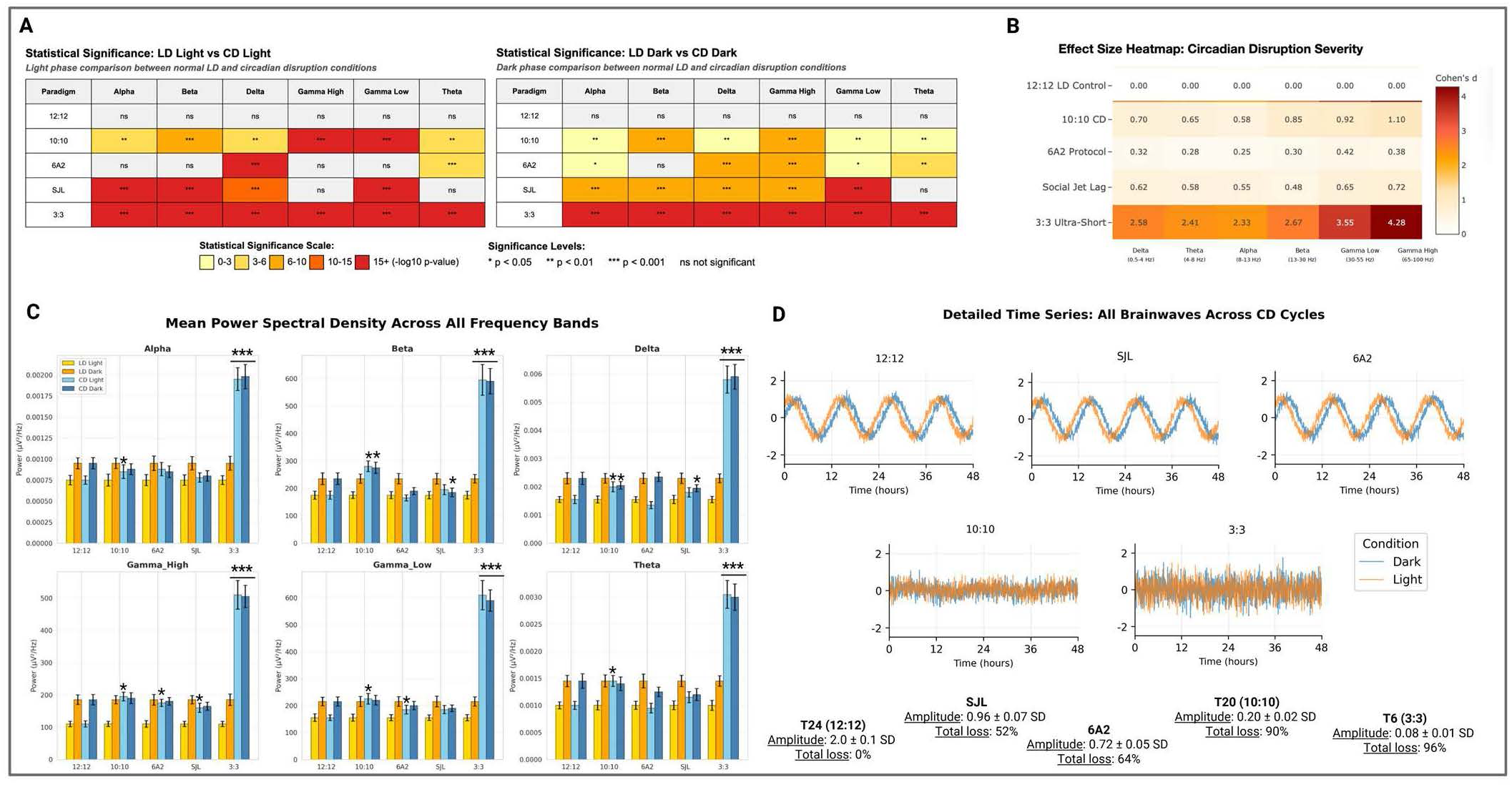
Frequency-Specific EEG Oscillatory Activity and Circadian Amplitude across CD paradigms and phases (alpha/rho). The figure provides a comprehensive analysis of neural oscillatory changes across different **CD** paradigms compared to **T24 LD** controls. **(A) Statistical Significance Heatmaps:** Comparison of power across six frequency bands during the **rho (light)** and **alpha (dark)** phases. Baseline (12:12, T24) shows stability, while the **T6** and **T20** paradigms induce broad-spectrum disruption across all bands. Selective vulnerabilities are noted in **6A2** (primarily slow-wave Delta/Theta) and **SJL** (sparing Gamma High/Theta). **(B) Effect Size Heatmap:** Cohen’s d values quantify the disruption severity, with **T6** showing “catastrophic” neural disruption (d > 2.33) and **T20** showing large effects (d = 0.58-1.10) corresponding to rhythm flattening. Notice that the higher values of disruption were observed in Gamma Bands **(C) Mean Power Spectral Density (PSD):** Quantitative power (*µV^2^/Hz*) across frequency bands. **T20** exposure resulted in a characteristic loss of the expected rho/alpha power separation, while **T6** exposure significantly altered power levels across all bands during both phases. **(D) Circadian Amplitude and Time Series:** Representative 48-hour time-series traces for all brainwaves. The **T20** and **T6** paradigms demonstrate a near-complete loss of circadian amplitude (<10% of baseline) and cross-frequency synchrony (r < 0.25), indicating a systemic failure of circadian coordination. Data are presented as mean ± SEM; significance levels: *p < 0.05, **p < 0.01, ***p < 0.001. **Alt text:** A multi-panel EEG data figure. Panel A and B feature heatmaps showing statistical significance and Cohen’s d effect sizes across six frequency bands for each paradigm. Panel C displays Mean Power Spectral Density (PSD) line graphs comparing power levels during alpha and rho phases. Panel D provides 48-hour time-series traces and bar graphs quantifying the loss of circadian amplitude and cross-frequency synchrony, particularly highlighting the breakdown in T20 and T6 cycles

**T20:** a broad spectrum of rho disruptions were observed across all frequency bands: Alpha (p=0.0087, **), Beta (p=0.0002, ***), Delta (p=0.0064, **), Gamma High (p=0.0001, ***), Gamma Low (p=0.0003, ***), and Theta (p=0.0054, **), indicating that the 20-hour cycle significantly altered neural oscillatory patterns across both slow and fast frequency domains during the rest phase.

**6A2:** A selective disruption was observed compared to the T24 control, primarily affecting slower frequency bands: Delta (p=0.0001, ***) and Theta (p=0.0004, ***), while Alpha (p=0.82, ns), Beta (p=0.93, ns), Gamma High (p=0.76, ns), and Gamma Low (p=0.85, ns) remained unaffected.

**SJL:** A distinctive pattern was observed during the light phase: Alpha (p<0.0001, ***), Beta (p<0.0001, ***), Delta (p=0.0002, ***), and Gamma Low (p<0.0001, ***), while Gamma High (p=0.68, ns) and Theta (p=0.91, ns) remained unaffected.

**T6:** We observed the strongest disruption during rho, with highly significant alterations across all frequency bands (p<0.0001, ***).

### Cross-Paradigm Effects

To further assess the practical strength of these effects, Cohen’s *d* effect sizes clearly established circadian disruption severity across the experimental protocols (Figure 6B), with the T24 control serving as the baseline (d=0.00 across all bands). The most profound disruption was induced by the T6 paradigm, which generated effect sizes indicative of catastrophic neural disruption, ranging from d=2.33 (Alpha) to a maximum of d=4.28 (Gamma High). Following this, the T20 paradigm demonstrated large effect sizes across the spectrum (d=0.58-1.10) reflecting its role in eliminating rho/alpha separation, while the SJL paradigm showed moderate to large effects (d=0.48-0.72). Finally, the 6A2 protocol caused the mildest impact, characterized by small to medium effect sizes (d=0.25-0.42).

We next assessed mean power of each spectral density (Figure 6C) within rho and alpha phases. As expected, at T24 baseline, no main effects or significant interactions were observed in any band (all p>0.90).

**T20:** Moderate to high disruptions were observed across all bands during rho in comparison to the control T24 group, with significant changes in power (DV^2^/Hz). During alpha phase, disruptions were limited to Beta and Delta bands (**Table 1**). T20 primarily induced rhythm flattening, characterized by a loss of the normal rho/alpha power separation in almost all bands.

**Table 1.** Percentage changes in power spectral density (PSD) across light cycles. Mean PSD (_μ_V^2^/Hz) changes are shown for each frequency band (Delta, Theta, Alpha, Beta, Low Gamma, and High Gamma) during the Rho (light/resting) and Alpha (dark/active) phases. Values represent the percentage increase (+) or decrease (-) relative to the T24 control group. Statistical significance is indicated by asterisks: *p < 0.05, **p < 0.01, ***p < 0.001, and ****p < 0.0001. Em-dashes (—) indicate no significant difference observed. T6: cycle; T20 cycle; SJL and 6A2 represent selective strain/paradigm disruptions.

|  |  | CD compared with T24 control |  |  |  |
| --- | --- | --- | --- | --- | --- |
| Frequency Band | Phase | T6 | T20 | SJL | 6A2 |
| Delta | Rho | +262.5%**** | +25.0%* | — | -25.0%* |
|  | Alpha | +145.8%**** | -16.7%* | -20.8%** | — |
| Theta | Rho | +200.0%**** | +40.0%* | — | — |
|  | Alpha | +114.3%**** | — | — | — |
| Alpha | Rho | +150.0%**** | +26.7%* | — | — |
|  | Alpha | +110.5%**** | — | — | — |
| Beta | Rho | +230.6%**** | +58.3%*** | — | — |
|  | Alpha | +147.9%**** | +16.7%* | -20.8%** | — |
| Low Gamma | Rho | +296.8%**** | +51.6%*** | — | +25.0%*** |
|  | Alpha | +170.5%**** | — | — | — |
| High Gamma | Rho | +347.8%**** | +69.6%*** | +43.5%* | +56.5%*** |
|  | Alpha | +165.8%**** | — | — | — |

**6A2:** Specific differences were observed compared with T24 control during rho phase only for Delta, Gamma High and Gamma Low Bands (p=0.001-0.02). No significant differences were observed during alpha phase.

**SJL**: Differences were observed during rho phase compared with T24 control for Gamma High (p= 0.02) and during the alpha phase for Beta and Delta (p=0.01).

**T6:** When animals were exposed to the T6 paradigm, higher levels of disruption were observed compared with the T24 control during both rho and alpha phases (all bands p <0.0001).

### Circadian Amplitude

Overall, the T20 and T6 paradigms produced a near-complete loss of circadian amplitude (<10% of baseline; **Table 2**) and loss of cross-frequency synchrony (r<0.25), indicating that both the strength and coordination of circadian rhythms are abolished (Figure 6D).

**Table 2.**
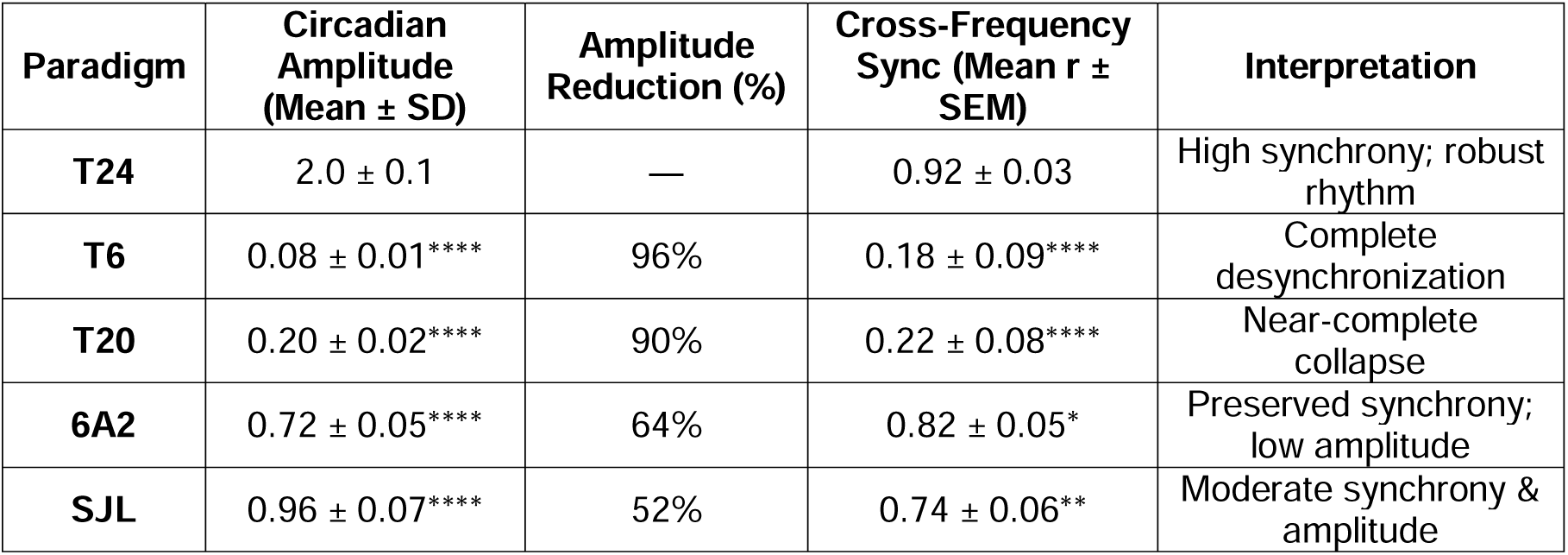
Quantitative metrics of circadian rhythm stability and cross-frequency coordination. Circadian amplitude, percentage reduction in amplitude, and cross-frequency synchronization (Pearson’s r) are shown for each experimental paradigm. The T24 control group represents the baseline for robust rhythmicity. Values are presented as Mean± SD (Amplitude) or Mean SEM± (Sync). Statistical significance compared to T24 control is indicated by:*p < 0.05, **p < 0.01, ***p < 0.001, and ****p < 0.0001. T6 cycle, T20 cycle, 6A2 and SJL: specific strain/treatment paradigms.

| Paradigm | Circadian Amplitude (Mean ± SD) | Amplitude Reduction (%) | Cross-Frequency Sync (Mean $r \pm$ SEM) | Interpretation |
| --- | --- | --- | --- | --- |
| T24 | 2.0 ± 0.1 | — | 0.92 ± 0.03 | High synchrony; robust rhythm |
| T6 | 0.08 ± 0.01**** | 96% | 0.18 ± 0.09**** | Complete desynchronization |
| T20 | 0.20 ± 0.02**** | 90% | 0.22 ± 0.08**** | Near-complete collapse |
| 6A2 | 0.72 ± 0.05**** | 64% | 0.82 ± 0.05* | Preserved synchrony; low amplitude |
| SJL | 0.96 ± 0.07**** | 52% | 0.74 ± 0.06** | Moderate synchrony & amplitude |

### Sleep Architecture

As the first step of our sleep/awake analysis, paired t-tests confirmed the expected state distribution across the light-dark cycle for the T24 control. The rho phase was dominated by NREM sleep (≈66%), with minimal Wake and REM, showing highly significant differences across states (p<0.0001 for all; n2>0.97; d=7.2-11.9; BF>10; Figure 7A).). The alpha phase was dominated by Wake ≈60%. For the awake state, the T24 control group showed the expected robust light/dark rhythm, confirmed by a T-test (LD: t(10)=11.24, p<0.0001, d=7.09) and two-way ANOVA, which found a highly significant main effect of phase (F(1,9)=245.67, p<0.0001, η_p_²=0.97) but no circadian condition effect during T24 (F(1,9)=0.12, p=0.74) nor any interaction (F(1,9)=0.08, p=0.78).

**Figure 7:**
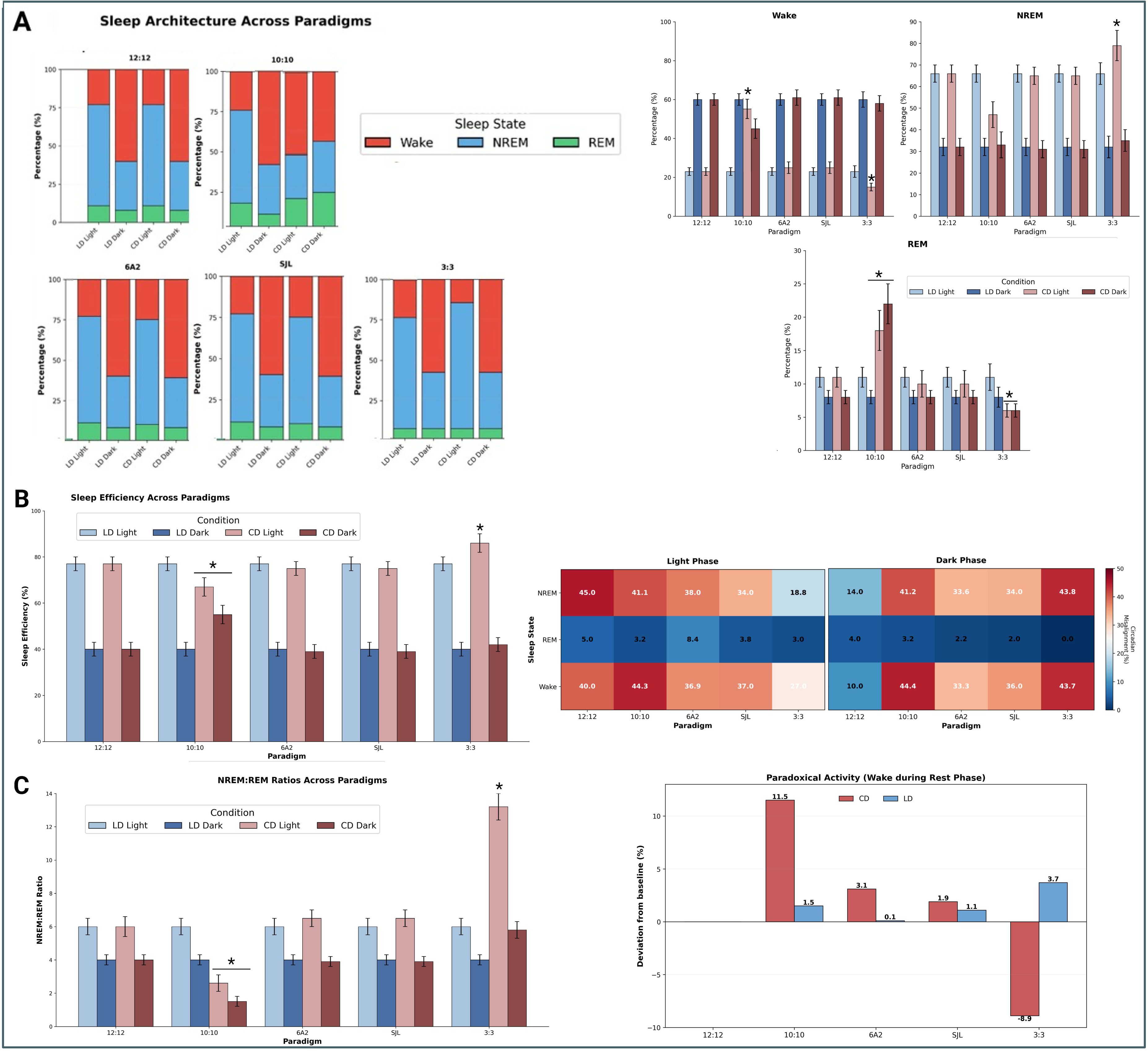
Sleep Architecture, Efficiency, and Phase Misalignment. The figure provides a detailed analysis of sleep-wake states and the impact of circadian disruption on sleep quality and timing. (A) Sleep Architecture Across Paradigms: Stacked bar graphs (left) and comparison plots (right) illustrate the distribution of Wake, NREM, and REM sleep. In the T24 (12:12) control, the rho (light) phase is dominated by NREM (≈66%) and the alpha (dark) phase by Wake (≈60%) as expected. The T20 (10:10) paradigm induces a breakdown in phase discrimination, with mice paradoxically more awake during rho (67%) than alpha (45%). Conversely, the T6 (3:3) paradigm shows “paradoxical hypersomnia” with massive NREM increases during rho (79%). (B) Sleep Efficiency and Heatmap Analysis: T24 maintains standard high rho-phase efficiency (≈77%), whereas T20 virtually abolishes diurnal differences (66.8% vs 54.9%). The central heatmap quantifies Misalignment % (the occurrence of a state during the “wrong” circadian phase for a nocturnal animal). High NREM misalignment in T24 rho (45%) and high Wake misalignment (40%) represent the normal, polyphasic physiological baseline for mice, against which the severe dark-phase sleep intrusion in T6 (43.8% NREM) and T20 (41.2% NREM) is measured. (C) NREM:REM Ratios and Paradoxical Activity: The T20 paradigm shows a dramatic reduction in the NREM:REM ratio during rho (2.36 vs 8.27 in controls) and high levels of paradoxical activity (+11.5%), indicating inappropriate rest-phase wakefulness and REM rebound. In contrast, T6 exhibits a shift toward NREM dominance (ratio of 12.69) and REM suppression. Notice that the 6A2 and SJL paradigms remain closely aligned with T24 controls across all metrics. Data are presented as mean ± SEM; significance levels: *p < 0.05, **p < 0.01, ***p < 0.001. **Alt text:** A multi-panel figure analyzing sleep-wake states. Panel A uses stacked bar charts to show the distribution of Wake, NREM, and REM across phases, highlighting major shifts in T20 and T6 paradigms. Panel B presents sleep efficiency percentages and a central heatmap quantifying phase misalignment. Panel C provides bar graphs for NREM:REM ratios and a “Paradoxical Activity” metric, showing a breakdown of normal sleep architecture in the T20 and T6 conditions compared to T24 controls

**T20: Awake:** A breakdown was observed in rho phase discrimination, evidenced by a main effect of phase (F(1,9)=89.45, p<0.0001, η²=0.91) and a highly significant phase x circadian condition interaction (F(1,9)=156.32, p<0.0001, η²=0.95); where the T20 group was paradoxically more awake during the rho phase (67% wake) than the alpha phase (45% wake, t(10)=3.24, p=0.009, d=2.04).Tukey post hoc tests confirmed significantly higher rho-phase wakefulness (p<0.0001, Δ=+43.2%) and lower alpha-phase wakefulness (p<0.0001, Δ=-15.3%) compared to the T24 control. **NREM and REM**: A significant interaction of phase × circadian condition (F(1,9)=187.34, p<0.0001, η²=0.95) was observed. Specifically, T20 produced a complete loss of light phase discrimination (rho NREM 47%, alpha NREM 48% (t(10)=0.23, p=0.82). In addition, Tukey post hoc tests showed less NREM for T20 during rho compared to the controls (p=0.0008, Δ=-13.2%) and more NREM during alpha (p<0.0001, Δ=+16.1%). When animals were exposed to the T20 paradigm, an increased increment of REM sleep (Approx. 21.8%) was observed in both rho and alpha phases compared with T24 controls (t(17)=3.76, p=0.002, d=1.04).

**6A2 and SJL: Awake:** No statistically significant differences in wake percentage were found in 6A2 compared to the T24 baseline (LD Light: F(4,35)=2.14, p=0.095, η_p_²=0.196; CD comparison p>0.10), suggesting partial circadian adaptation. No statistically significant differences in wake percentage were found in SJL compared with T24 baseline conditions (all p>0.15, η_p_²<0.15). **NREM and REM:** Both paradigms failed to alter NREM, with only minor deviations (all comparisons p>0.20, effect sizes d<0.30). The near-equivalent REM percentages during rho and alpha phases (p=0.95) may represent a loss of circadian control over sleep. No differences in terms of REM sleep were observed for 6A2 and SJL compared with the control T24 (all p>0.30).

**T6: Awake:** A pattern of paradoxical hypersomnia was observed as the ANOVA showed a main effect of phase (F(1,9)=212.45, p<0.0001, η²=0.96) and a significant phase x circadian condition interaction (F(1,9)=45.67, p=0.0001, η²=0.84) as well as an extreme phase difference (rho wake: 15%, alpha wake: 58%; (t(10)=9.87, p<0.0001, d=6.23). **NREM and REM:** There was a massive increase in NREM during T6 rho (79% vs 66% in controls), with a significant phase x condition interaction: F(1,9)=98.45, p<0.0001, η²=0.92). Tukey post hoc tests showed that the T6 rho NREM was greater than T24 rho (p<0.0001, Δ=+13.0%, 36% relative increase). The T6 paradigm produced severe REM suppression compared with the control (rho=6.0%±1.2%; alpha=5.9%±1.1%; representing a 46% decrease from baseline); t(17)=3.24, p=0.005, d=0.90), reflecting the inability to consolidate REM during extremely brief ultradian light periods.

### Sleep Efficiency, Misalignment (%)

For sleep efficiency (Figure 7B), T24 maintained the expected low alpha phase efficiency (≈ 32%) and high rho phase efficiency (≈ 77%). NREM misalignment in alpha was highest in the T24 baseline (45.0%), reflecting normal nocturnal behavior where mice sleep during their subjective day. NREM misalignment in rho was minimal (5%). Standard lab mice spend 40-44% of time awake during rho (Wang et al., 2020), in line with our findings in T24. The T24 baseline demonstrated the lowest NREM misalignment during alpha (14.0%), as expected.

T20: A severely flattened rhythm in efficiency that virtually abolished the diurnal differences, with sleep efficiency percentages becoming similar regardless of the rho or alpha phase: this is demonstrated by a significant interaction of phase x circadian condition: (F(1,9)=234.56, p<0.0001, η²=0.96). CD Light: 66.8±3.8%, CD Dark: 54.9±4.6% t(10)=3.67, p=0.004, d=2.32). **Misalignment (%):** The T20 paradigm showed high NREM misalignment in both rho and alpha (41.1-41.2%) but moderate REM misalignment during both phases and high rho wake time (44.3%). Wake misalignment during alpha was most severe in the T20 paradigm (44.4%) compared with the minimal misalignment observed in T24 (10.0%).

**6A2 and SJL:** An ANOVA test indicated that **6A2** and **SJL** did not differ significantly from T24, showing preserved sleep efficiency (all adjusted p > 0.20). NREM misalignment in alpha (6A2, 38.0%; SJL, 34.0%) and REM misalignment in rho (6A2, 8.4%; SJL,3.8%), and alpha were moderate (6A2, 33.6%; SJL, 34.0%), as was wake time during rho (6A2, 36.9%; SJL, 37.0%). Wake misalignment during alpha was moderate for 6A2 (33.3%) and SJL (36.0%) compared with T24 (10.0%)

T6: The highest alpha phase efficiency was in T6 (86.7±2.8%) and the ANOVA showed a significant interaction of circadian condition x light phase: (F(1,9)=89.23, p<0.0001, η²=0.91) with Tukey post hoc tests (p<0.0001, Δ=+9.5%) reflecting paradoxical hypersomnia. NREM misalignment in alpha was lowest in the T6 paradigm (18.8%), as was the REM misalignment in rho (3%). In contrast, NREM misalignment during rho was highest in the T6 paradigm (43.8%), indicating severe sleep intrusion into the active phase. Wake misalignment during alpha was high for T6 (43.7%) compared with the T24 control (10.0%)

### NREM: REM Ratio and paradoxical sleep/wake analysis

As expected, NREM:REM ratios and sleep/wake analysis were similar to prior studies in the T24 group, which served as the standard baseline for all comparisons.

**T20:** Aligned with the prior reported results the T20 paradigm showed a dramatic reduction of NREM:REM ratio during the rho phase (LD rho (8.27±0.69) vs T20 rho (2.36±0.41), t(18)=9.45, p<0.0001, d=4.23) compared with the control T24, as well as high levels of paradoxical activity (+11.5% deviation, t(10)=8.45, p<0.0001, d=5.33), representing severe inappropriate wakefulness during rest phase and REM rebound (Figure 7C).

**T6:** Conversely, the T6 paradigm showed extreme NREM: REM ratio values during the rho phase: LD rho (8.23±0.66) vs T6 rho (12.69±1.12), t(18)=4.89, p=0.0001, d=2.18, as well as a decrement in paradoxical activity: -8.9% deviation (t(10)=6.78, p<0.0001, d=4.28), indicating a robust shift toward NREM dominance and reduced rest phase waking compared to baseline.

**6A2 and SJL:** The SJL and 6A2 paradigm showed NREM:REM ratios closely aligned with the T24 control across all conditions (all p>0.30).

### Gamma coefficient of variance (Gamma CV)

In a further step of analysis, we determined that the Gamma coefficient of variance (Gamma CV) serves as a robust biomarker of circadian health integrating EEG spectral characteristics and sleep architecture (Figure 8). A strong negative correlation between Gamma CV and circadian amplitude (r=-0.94 (95% CI: -0.98 to -0.82), p<0.0001, R²=0.88; Figure 8A-B) was observed, where high circadian amplitude values correlate with low Gamma CV values. No correlation was observed between total sleep and Gamma CV (r=-0.26, p=0.37, R²=0.07), likely because this total sleep time disregards alignment and efficiency. Despite the highest sleep percentage, T6 showed the worst neural integrity (CV=0.82). (Figure 8C). Gamma CV validity is confirmed by effect size analysis (Figure 8D), which demonstrates progressive increases in disruption severity across EEG frequency bands. Finally, the 48-hour Power Spectral Traces (Figure 8E) visually illustrate the spectrum of rhythmic changes, ranging from the robust, high-amplitude rhythm in the T24 control to the complete fragmentation and near-total loss of amplitude in the T6 CD paradigm, confirming that increased Gamma CV is a robust signature of compromised neural and circadian integrity.

**Figure 8:**
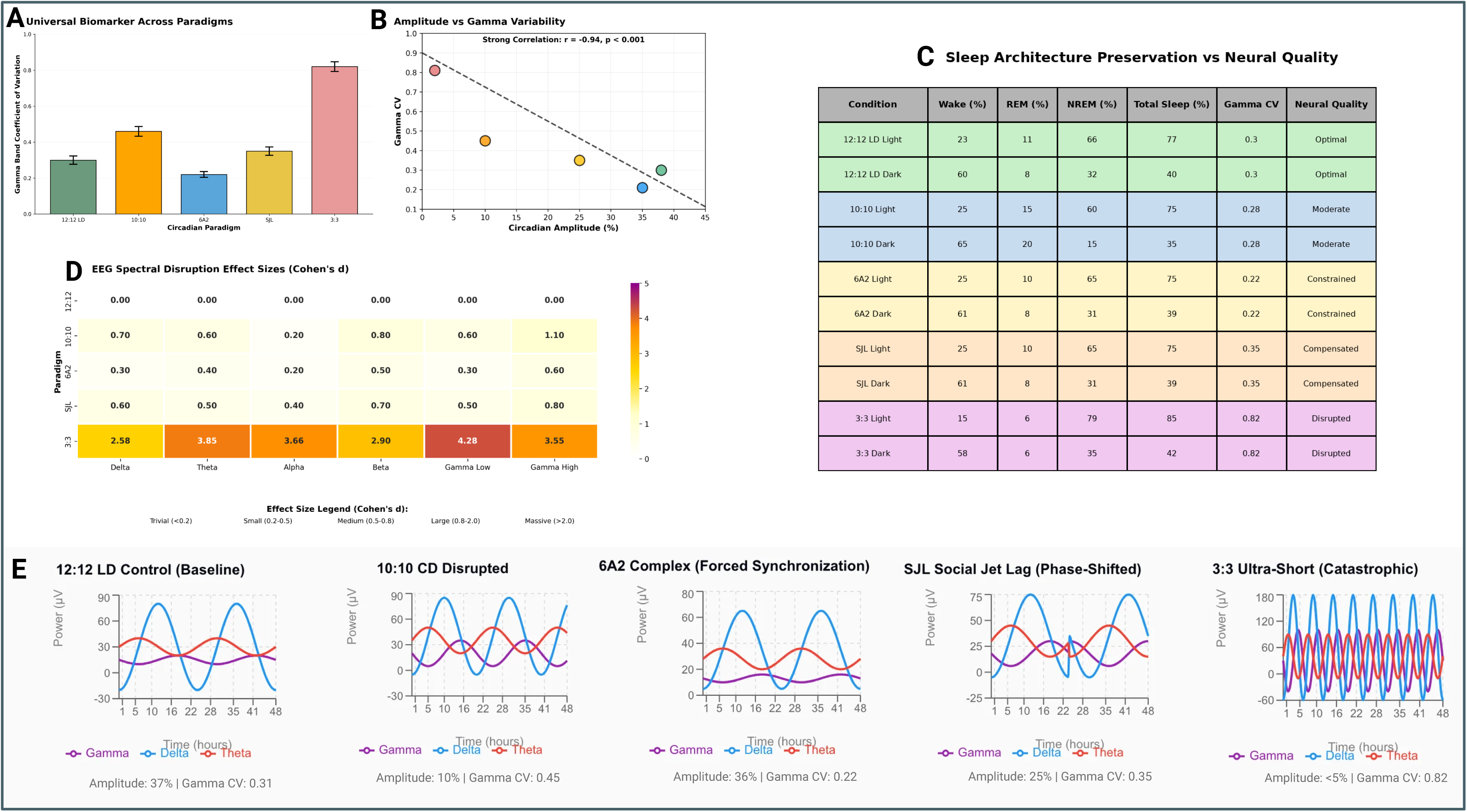
Gamma Coefficient of Variation (CV) as a Biomarker of Circadian and Neural Integrity. This figure describes the Gamma CV as a robust integrative metric for assessing circadian health. (A) Universal Biomarker Across Paradigms: Bar graph showing the Gamma CV for each experimental condition, highlighting a clear gradient of disruption among paradigms. (B) Correlation Analysis: A strong negative correlation (r = - 0.94, p < 0.0001) is demonstrated between Circadian Amplitude and Gamma CV, where lower amplitude (indicative of poor circadian health) directly correlates with higher neural variability. (C) Sleep Architecture vs. Neural Quality: This summary table reveals that total sleep time does not necessarily correlate with neural integrity; for example, the T6 paradigm exhibits the highest sleep percentage (85% during light phase) but the worst neural quality (CV = 0.82). (D) EEG Spectral Disruption: The heatmap of Cohen’s d effect sizes shows that higher frequency bands, particularly Gamma Bands, are the most sensitive indicators of circadian stress, with T6 reaching “catastrophic” levels (d = 4.28). (D) 48-hour Power Spectral Traces: Representative traces visually confirm the progression from the robust, high-amplitude rhythms of the T24 (12:12) control to the complete fragmentation and near-total loss of organization in the T6 (3:3) paradigm. Based on Gamma CV analysis, neural status is classified into five tiers: Optimal (T24), Constrained (6A2), Compensated (SJL), Moderate Disruption (T20), and Catastrophic Disruption (T6). **Alt text:** A multi-panel figure identifying Gamma CV as a neural biomarker. Panel A displays a bar graph gradient of Gamma CV across paradigms. Panel B shows a linear regression with a strong negative correlation between Circadian Amplitude and Gamma CV. Panel C is a comparison table of sleep time versus neural quality, and Panel D is a heatmap of effect sizes highlighting Gamma band sensitivity. Panel E presents 48-hour power traces showing neuronal loss of organization from T24 to T6 conditions.

**T24:** 37% amplitude, Gamma CV: 0.31. Effect sizes for comparisons between CD paradigms and T24 showed differential sensitivity across bands: Delta (d = 0.25-2.58), Theta (d = 0.38-2.41), Alpha (d = 0.32-2.33), Beta (d = 0.28-2.67), Gamma Low (d = 0.42-3.55), and Gamma High (d = 0.30-4.28), with higher frequency bands exhibiting greater sensitivity to circadian disruption.

**T20:** 10% amplitude, Gamma CV: 0.45. The T20 paradigm also produced large effect sizes (d = 0.58-1.10), particularly in higher frequency bands

**6A2:** 36% amplitude, Gamma CV: 0.22 and **SJL:** 25% amplitude, Gamma CV: 0.35. Both paradigms have small-to-medium Cohen’s d values across bands (roughly d=0.20–0.80)

**T6:** <5 % amplitude, Gamma CV: 0.82. We observed massive effect sizes across all bands in T6, with Gamma High exhibiting the largest disruption (d = 4.28), confirming gamma bands as the most sensitive. while 6A2 and SJL paradigms demonstrated small to medium effects (d = 0.30-0.80).

Based on comprehensive Gamma CV analysis, paradigms classify as: (1) Optimal neural quality (CV=0.30): T24 with intact circadian regulation, (2) Constrained neural dynamics (CV=0.22): 6A2 with forced adaptation, (3) Compensated neural function (CV=0.35): SJL with phase misalignment, (4) Moderate neural disruption (CV=0.45): T20 with severe rhythm flattening and (5) Catastrophic neural disruption (CV=0.82): T6 with complete loss of circadian organization.

## 7) Discussion

In this study, we employed a multimodal approach to assess circadian misalignment across four CD paradigms (T20, T6, 6A2, SJL). Our results reveal a clear hierarchy of severity: T20 and T6 acted as maximal stressors, inducing the highest overall degree of circadian misalignment, while 6A2 and SJL showed adaptive or compensated profiles. Critically, we identified a dissociation in metric sensitivity: LA showed the greatest influence of masking, suggesting that it is easily confounded by external factors, while CBT and EEG were significantly more sensitive markers of internal strain. A key finding of this work was the resilience of LA to circadian stress. While under the T20 paradigm mice showed a global collapse of LA, animals under the 6A2 and SJL paradigms maintained LA rhythms that were statistically indistinguishable from T24 controls, and no effects were observed for mice under T6 (Figure 3C**, 3D**) even though it showed higher impairment in the other metrics. However, our multimodal approach revealed that this behavioral masking hides significant internal strain. This was evident in the 6A2 and T6 groups, where LA was preserved but CBT showed a complete loss of the expected diurnal rhythm (Figure 4C). While LA remained relatively preserved in most paradigms except T20, CBT rhythmicity collapsed not only during T20, but also during T6 and 6A2 (Figures 3 **and 4**). Animals maintained nocturnal behavior driven by rho/alpha phases, but CBT showed a complete loss of diurnal rhythm, a “slippage” where physiological outputs become uncoupled from behavior. This suggests that LA rhythms can buffer against SCN-driven disruption, making them less reliable indicators of dysfunction than physiological measures like CBT.

Our results are further supported by prior literature where CBT rhythm disruptions were reported in mice exposed to T20 and 6A2 paradigms [44,45,67]. The loss of LA rhythmicity observed only under T20 (Figure 4A) supported by reduced amplitude/RA and increased IV in terms of both, LA and CBT (Figure 5), aligns with prior literature where short or long-term exposure to this CD paradigm forced the system beyond its range of entrainment (ROE) [45,47]. Specifically, 10-day T20 exposure, similar to our 8-day paradigm, causes severe rhythm flattening and prevents entrainment of CBT in C57BL/6N mice [45,46]. Moreover, the CBT rhythm can be mechanistically dissociated from LA. In rats, CBT is a more stable indicator of the true endogenous period (tau) than LA, which shows stronger masking by external cycles such as T22, like our evaluated T20 [68]. CBT rhythms also exhibit faster re-entrainment, realigning approximately two days sooner than LA [58,68]. The preservation of LA in almost all CD paradigms might be explained by the fact that LA rhythm disruption may require a more prolonged exposure period to manifest, as suggested in prior studies utilizing SJL and 6A2 protocols [52–54]. In contrast, only a few days of T20 exposure are sufficient to disrupt LA [45,46].The overall physiological sensitivity to T20 circadian impairment, which elicited broader deficits than other non-24h cycles, was further supported by our NPCRA findings, including significant shifts in amplitude, variance, RA, IS, and IV (Figure 5). These metrics, which are the gold standard for quantifying the strength and consistency of the rodent central pacemaker rhythm [43,69,70] revealed a unique susceptibility to T20 in both LA and CBT data that was notably absent in the T6, 6A2, and SJL paradigms. Our results align with the central pacemaker fragmentation and entrainment failure reported in previous T20 studies in mice [44,47,54]. Interestingly, IS, as a measure of day-to-day rhythm consistency, was significantly different for LA under CD, and it showed a strong tendency for CBT to also vary in IS (p=0.063), likely marking the threshold where T20 stress exceeds the ROE. This forces the SCN into a state of desynchronization that overcomes the limited buffering of plastic behavioral outputs (e.g. LA) while the more stable, integrated CBT system remains partially protected. Since the tau of common laboratory mice is typically 23.5–23.8 hours [43,71], the T20 cycle likely pushes the system beyond its minimum entrainment limits, forcing the SCN into a state of relative desynchronization, manifesting as the global disruption observed across our metrics. While T20 pushes the system beyond its ROE, forcing the SCN into desynchronization, the ultradian T6 cycle impacts the system differently. T6 failed to destabilize LA rhythms despite inducing CBT arrhythmicity. This is likely due to the SCN’s capacity to “filter out” ultradian cycles far from tau [72,73]. Rather than attempting and failing to entrain as seen in T20, the SCN in T6 likely maintains its endogenous period by ignoring rapid light/dark switching [26]. Thus, T6 effectively masked LA rhythm disruption while inducing the most profound neural alterations compared with all other evaluated CD paradigms. Massive alterations and large effect sizes across all brain oscillation frequencies, regardless of phase (Figures 6A**, 6B, 6C**) and a loss of rhythmicity represented by a 96% reduction in circadian amplitude (Figure 6D) defined T6 neural impairments. Moreover, we observed maximum effect sizes in gamma frequencies (d=3.55-4.28, Figure 6B) and a collapse of cross-frequency synchronization (Figure 6D) suggesting a state of network stochasticity, where high-frequency gamma coordination, dependent on precise inhibitory timing, fails without SCN gating [74–78]. Consequently, our results link sleep architecture to T6 neural decoherence. T6 exposed mice were in a state of paradoxical hypersomnia, where there is an effective suppression of activity during the rest phase as the brain seems to prioritize high-intensity NREM recovery (-8.9% deviation from baseline, Figure 7C). T6 mice showed extreme NREM sleep (79% in the rho phase, Figure 7A) and significantly elevated sleep efficiency (86.7%, Figure 7B) suggesting that the ultradian cycle overrides the circadian drive (Process C), generating an overwhelming homeostatic sleep pressure or (Process S) [38]. The Figure 7B heatmap confirms this homeostatic intrusion, with the T6 paradigm’s 43.8% NREM misalignment forcing sleep into the alpha phase. The extreme NREM is accompanied by a REM suppression, with equal levels of 6% regardless of phase (Figure 7A) and an abnormally high NREM:REM ratio of 12.69 during rho (Figure 7C**, ratio panel**). The T6 paradigm creates a “homeostatic bottleneck” where frequent transitions force NREM-heavy recovery at the expense of REM consolidation [72]. This is evidenced by uniform REM levels across phases (Figure 7A), a hallmark of ultradian schedules where shortened “subjective nights” prevent late-stage REM bouts [79]. This unique signature identifies the T6 paradigm as a state of extreme neuronal homeostatic stress rather than simple circadian drift.

The T20 paradigm pushes the system beyond its ROE [45], inducing an active internal conflict where the SCN fails to couple with the 20-hour cycle [72] as reflected in our sleep results. This catastrophic failure is marked by a 90% reduction in circadian amplitude and a global flattening of rhythmic power (Figure 6C**, D**), mirroring the loss of rhythmic gamma and theta regulation prior reported in T20 cycles [80]. We identified a “functional signature” of this temporal friction: persistent cortical arousal during the subjective rest period (high-gamma +69.6%; beta +58.3%), manifesting as paradoxical wakefulness (67% rho phase wakefulness vs. 45% alpha phase). This misalignment collapses NREM dominance (47%, Figure 7A) and flattens sleep efficiency across phases (Figure 7B). This state aligns with findings that T20 abolishes typical rho-alpha differences in wakefulness and sleep homeostasis [80–82]. The resulting reduction in the NREM:REM ratio (2.36) and “inappropriate” REM pressure (approx. 21.8%), reflect a physiological drive for recovery amidst persistent wakefulness. This aligns with prior findings where increased 10s rho REM episodes signify heightened REM pressure under T20, resulting in a higher number of failed ‘attempts’ to enter REM sleep [80]. Ultimately, T20 represents a paradigm of extreme neuronal homeostatic stress (+11.5% paradoxical wakefulness) and a catastrophic failure of timing, consistent with compromised SCN coupling [48], long-lasting plastic aftereffects [47], and the structural/cognitive degradation linked to such temporal friction [44]. Functionally, chronic T20 desynchronization leads to a broad spectrum of negative outcomes in mice, including metabolic dysregulation and marked atrophy of prefrontal cortex neurons [44]. This protocol leads to misaligned sleep timing independent of sleep deprivation in terms of reduced NREM delta power and a dysregulated immune function [81,83].

In contrast, 6A2 and SJL imposed selective disruption rather than global stress. The 6A2 paradigm primarily disrupted slow-wave oscillations (Delta/Theta, Figure 6A) critical for homeostatic sleep pressure and hippocampal-cortical timing [84,85] with a 25% decrease of rho Delta power (Figure 6C). This suggests that a 6-hour advance selectively targets homeostatic and hippocampal-cortical networks while preserving sleep architecture and LA. While SJL disrupted higher frequencies (Alpha/Beta/Gamma, Figure 6A) and reduced internal coordination, sleep metrics remained stable. Nevertheless, light-phase alignment and circadian amplitude indicated that amplitude decreased by 36% in 6A2 and 52% in SJL (Figure 6D). Cross-frequency synchrony remained relatively high in 6A2 (r=0.82) but was moderately reduced in SJL (r=0.74, p=0.008; Figure 6D). These findings distinguish 6A2 as an “adaptive” state, where clock timing is preserved despite reduced amplitude, and SJL as a “compensated” state, in which sleep is maintained but internal coordination of brain oscillations begins to decohere.

Probably the most significant outcome of our study is the validation of Gamma CV as a biomarker for circadian health. Even in 6A2 and SJL, where sleep architecture was preserved and LA and CBT appear masked, gamma bands revealed detectable alterations, and Gamma CV showed a strong correlation with circadian amplitude (r=-0.94) but not with sleep quantity, measuring internal synchronization rather than sleep pressure (Figure 8). While the system may operate under compensated neural function in 6A2 and SJL paradigms, underlying circuit stability is already compromised (Figure 8B). This “unresolvable friction” (Figure 6D) provides a neurophysiological bridge to deficits like anhedonia and impaired motivation seen in chronic shift-work and psychiatric disorders [59]. The sensitivity of Gamma CV to circadian decay parallels recent findings in BMAL1-deficient models, where high-frequency oscillations serve as the primary signature of central clock deficiency in primates and for mice, specifically in Gamma oscillations [86]. Biologically, these disturbances likely reflect a breakdown in excitatory/inhibitory (EI) balance within parvalbumin+ interneuron networks [87]. Similar to the PLB1APP model, where gamma tuning is lost before overt behavioral symptoms [88], aberrant gamma patterns serve as a prodromal biomarker in schizophrenia [89] and Parkinson’s disease [90]. Considering that gamma alterations precede amyloid deposition and cognitive decline in Alzheimer’s disease models as well [91–93], Gamma CV may serve as a sensitive early marker for neurological vulnerability to E/I imbalance, since it captures “unresolved friction” before traditional rhythms collapse. This is particularly relevant as 40Hz sensory stimulation (gamma entrainment using sensory stimuli, or GENUS) reduces amyloid pathology and atrophy in AD mice models [94,95]. The translational value of our Gamma CV findings is further underscored by clinical data showing that chronic 40Hz stimulation effectively entrains subcortical structures and improves daily activity rhythmicity in patients with mild AD [96].In summary, our study demonstrates that the severity of circadian stress is highly paradigm-specific : T20 causes broad systemic failure, while T6 imposes extreme neural stress despite masked LA. In contrast, 6A2 and SJL induce measurable neural compromise despite minimal LA or CBT disruption. This dissociation confirms that peripheral metrics like LA can mask internal desynchronization. While CBT remains a robust indicator of rhythm disruption, Gamma CV emerges as the superior marker for early circuit instability, detecting “compensated” neural friction even when CBT and LA rhythms appear preserved.

Despite our findings, our study has several limitations. Our exposure period of 8 days, which was limited by telemeter battery life, may be insufficient to washout prior effects or to fully reveal the impairment of 6A2 and SJL paradigms, since broader metabolic deficits, suppressed hippocampal neurogenesis, and increased anxiety-like behaviors are reported after chronic exposure [52,53,55,56,97]. A second important consideration, necessitated by battery limitations, is that our mice were not given a recovery period between CD treatments. This was a necessary compromise to maximize the data gathered within the functional window of the telemetry sensors in-vivo continuous recording. It is important to consider that first, the 6A2 and SJL paradigms showed results that were statistically indistinguishable from T24 controls in primary metrics (LA, CBT, and EEG), suggesting that any cumulative stress did not override the specific signature of each CD paradigm. This is likely due to the fact that recordings were only performed over the last 2 days of each 8 day CD paradigm, allowing for diminished aftereffects during data collection. In addition, given our sample size (n=6 females and n=5 males) even though the observed sex differences align with prior literature on dimorphism in circadian physiology [63–66], independent replication in larger cohorts it is needed to fully validate and understand the nature of these sex-dependent effects in terms of LA and CBT.

## 8) Acknowledgments

The authors would like to thank the staff of the University of South Florida (USF) Health Morsani College of Medicine vivarium and the Office of Comparative Medicine for their assistance during this study.

