## Supplementary material for "Gamma CV as a Marker of Circadian Disruption in C57BL/6J Mice: Correlating Neural Desynchrony with Locomotor, Thermal, and Sleep Dysrhythmia across a Spectrum of Circadian Rhythms Disruption paradigms": Figures S1 and S2

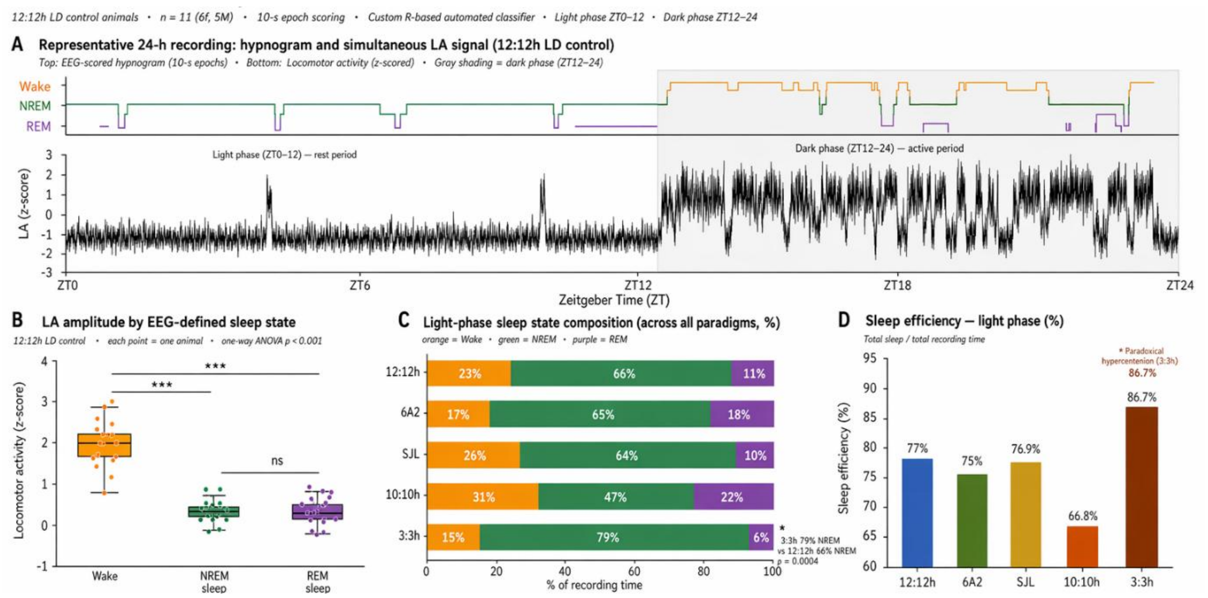


**Supplementary Figure 1. Validation of locomotor activity (LA) signal as a proxy for automated EMG-based sleep-state classification.**

**(A) Representative 24-h recording from a single 12:12h LD control animal.** Top panel displays the EEG-scored hypnogram at 10-s epoch resolution, identifying Wake (orange), NREM (green), and REM (purple) states. Bottom panel shows the simultaneously recorded LA signal (z-scored). The light phase (ZT0–12) is the primary rest period, characterized by consolidated NREM, periodic REM bouts (~1.1%), and low LA amplitude, while the dark phase (ZT12–24) represents the active period. Gray shading indicates the dark phase.

**(B) Boxplots of mean LA amplitude (z-scored) by EEG-defined sleep state.** Data represents mean values in in animals during the 12:12 T24 cycle (n = 11; 6F, 5M). Each point represents an individual animal. *** p < 0.001; ns, not significant (one-way ANOVA). LA amplitude is significantly higher during Wake compared to NREM and REM sleep.

**(C) Light-phase sleep state composition across all paradigms (%).** Stacked bar charts show the percentage of recording time spent in Wake (orange), NREM (green), and REM (purple) for different light cycles (12:12h, 6A2, SJL, 10:10h, and 3:3h). The 3:3h paradigm shows a significant increase in NREM (79%) compared to the 12:12h control (66%; p = 0.0004).

**(D) Sleep efficiency during the light phase (%) across paradigms.** Sleep efficiency is calculated as (total sleep / total recording time) during the light phase. Animals maintained in an ultra-short 3:3h cycle exhibit significantly higher sleep efficiency (86.7%; $p = 0.0004$) and paradoxical hyper-retention.

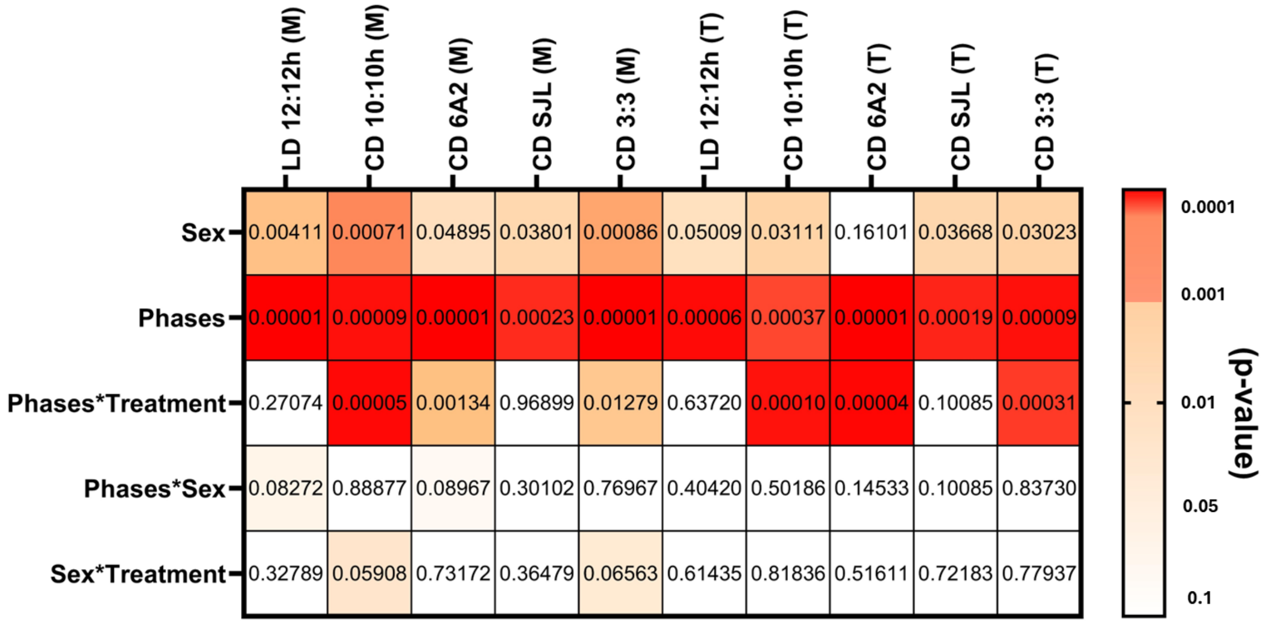


**Supplementary Figure 2: Statistical Analysis of Sex-Dependent Effects.** Heatmap displaying p-values for the main effects and interactions of sex across experimental paradigms (M: LA; T: CBT). A significant main effect of sex was observed for both LA and CBT ($p < 0.0001$ to $p < 0.05$; power 0.53-0.99), with females exhibiting higher levels than males. This effect was consistent across all comparisons and showed no significant interaction with circadian paradigm (LD or CD) or phase (alpha or rho). Crucially, the absence of significant sex x condition interactions confirms that sexual dimorphism did not account for the primary effects of the circadian paradigms on LA, CBT. No sex effects or interactions were observed for power spectrum density, sleep, Gamma CV, or EEG analysis.
